# Robust ORF Deconvolution from Multiple Ribo-seq Datasets with Group LASSO Penalty

**DOI:** 10.64898/2026.10.01.755973

**Authors:** Kilian Maidhof, Andreas Schlosser, Florian Erhard

## Abstract

Ribo-seq has provided compelling evidence that there are many translated but unannotated ORFs. Despite a growing body of peptidomic support for cryptic ORFs, the computational discovery of weakly translated, short and mutually overlapping ORFs has remained a major bottleneck. Here we introduce PRICE 2, a method designed to resolve these challenging translation events by adapting concepts from isoform deconvolution. PRICE 2 employs a generative model combined with group LASSO regularized regression to jointly infer active ORFs across multiple sequencing runs. In a comprehensive benchmark, PRICE 2 out-performed thirteen state-of-the-art tools in detecting short and overlapping ORFs. Orthogonal validation via immunopeptidomics demonstrated that ORFs identified by PRICE 2 enable the discovery of ∼40% more cryptic MHC-I peptides than the best-performing competing tool at a fixed false discovery rate. By analyzing 180 public ribo-seq samples spanning 45 cell and tissue types, we constructed *ORF-stock*, a comprehensive database of non-canonical ORFs for immunopeptidomic studies.

## Introduction

Eukaryotes typically encode exactly one protein per mRNA from a single canonical open reading frame (ORF; here, we use the term ORF for a translated region on a transcript from a start to a stop codon). However, translation outside canonical ORFs (cORFs) is pervasive [1] and has been described in many distantly related eukaryotic taxa (e.g., yeast [2], plants [3], and humans [1]). Whether it is mostly translational noise or serves a purpose is largely unknown. Some non-canonical ORF (ncORF) products have a demonstrated function [4]. Non-canonical translation events themselves can also be regulatory, such as upstream ORFs that inhibit their downstream cORF [5]. Some ncORFs can also evolve a function, suggesting that their high frequency is a source of newly evolving proteins [6]. Recent structured collaborative efforts have annotated functional microproteins and short ORFs that are translated without evidence for a functional product [7].

Translation of ncORFs can also have an important role in immunosurveillance. The proteasome breaks down proteins at the end of their lifespan and defective ribosomal products into shorter peptides [8]. The major histocompatibility complex I (MHC-I) presents peptides of roughly 8-12 aa at the cell surface. CD8+ T-cells can recognize mutated and virus-infected cells via aberrant peptides. Immunopeptidomics, which determines the set of presented peptides, has shown that 2-10% of them cannot be explained by cORFs but by translated ncORFs [9]. The fraction of detectable cryptic peptides depends on the MHC-I type and increases with virus infection and stimulation with pro-inflammatory cytokines [9, 10], suggesting an important role of non-canonical translation in immunosurveillance. Furthermore, for some types of cancer most neo-antigens stem from ncORFs, making the detection of cryptic peptides a viable approach to identify drug targets [11]. Immunopeptidomics data are also suitable as orthogonal evidence to benchmark ORF predictions from ribo-seq data [12]. The signals used to detect ORFs include evolutionary conservation, lack of stop codons in long sequence intervals, peptide sequence evidence from mass spectrometry data, and evidence from ribo-seq experiments. Detecting ncORFs from these signals is challenging because they are less conserved, shorter, often lowly translated, and their peptide products can be short-lived and therefore not detectable in mass spectrometry experiments [13]. Many current efforts to overcome these challenges focus on ribo-seq data. Ribo-seq, or ribosome profiling, is a sequencing approach that produces reads of ribosome-protected fragments (RPF), i.e., RNA covered by ribosomes [14]. These RPFs are mostly located within ORFs. The read start coverage shows a characteristic 3 nt periodicity that reflects the translocation steps of the ribosome and the precise cuts that RNase I, typically used for ribo-seq, makes at the boundary of the ribosome. Ribo-seq data also show higher coverage at translation start and stop sites. Translation initiation inhibitors such as harringtonine or lactimidomycin can further enrich the start site signal [15].

Several computational tools detect ORFs from ribo-seq data [12, 16–18]. Most first generate a set of ORF candidates and then determine which of them are active. One general approach, applied in different variants, computes various metrics for all ORF candidates and then applies thresholds or learns decision boundaries with machine learning models to distinguish active from inactive ORFs. Important and frequently applied metrics include measures for periodicity and increased coverage at translation start and stop sites [19]. PRICE 1 takes another approach and deconvolves the read signal to estimate an activity at the codon level. It then stitches codons together to find a parsimonious set of ORFs and employs a stringent statistical test to distinguish signal from noise [12]. Although existing tools perform well at detecting cORFs, detecting ncORFs remains challenging. Low coverage, short length, and overlap with other ORFs complicate this task, especially for internal ORFs (iORFs; ORFs completely contained within a cORF), where the overlapping ORF masks the periodicity signal [20].

Here, we introduce PRICE 2, a tool that overcomes these limitations through four major improvements to earlier methods: (i) PRICE 2 implements a generative model of read distributions across ORFs that accounts for the 3nt periodicity and uneven coverage across ORFs; (ii) Extending the classical isoform deconvolution model for RNA-seq data [21], PRICE 2 distributes reads across overlapping ORFs according to this generative model. The redistribution considers that reads may map to multiple locations; (iii) It applies a group LASSO penalty to share information on translated ORFs across multiple samples to improve sensitivity; (iv) It applies a robust Huber Poisson regression to mitigate the effects of uneven read coverage over ORFs inherent to ribo-seq data.

We show that in simulated data PRICE 2 has higher sensitivity for more active ORFs and for larger numbers of samples, and that this gain covers all types of ORFs. Comparing ORF discovery of most existing tools using immunopeptidomics evidence at a fixed false discovery rate (FDR), we demonstrate that the gain in sensitivity does not sacrifice specificity. At a fixed FDR, PRICE 2 improved sensitivity for the identification of cryptic peptides by 40%. We also introduce *ORF-stock*, a comprehensive database of cryptic ORFs generated by applying PRICE 2 to 180 publicly available ribo-seq samples selected for high quality and diversity in genetic activity.

## Results

The statistical model behind PRICE 2 draws on the codon level model of PRICE 1 and on the well-established generative model for isoform deconvolution from RNA-seq data [21]. In isoform deconvolution, short RNA-seq reads are generated from mutually overlapping transcripts with probabilities depending on their length and abundance. Because of these overlaps and the short read length, many reads could originate from multiple isoforms. The transcript origin is therefore treated as a latent variable, and isoform abundances are estimated with EM algorithms, other maximum likelihood or variational Bayes approaches.

This idea extends to ORF detection and quantification when the model accounts for additional ribo-seq specific signals (Fig. 1). PRICE 2 integrates conditional probabilities for periodicity in the three phases relative to the ORF the reads originate from, for stronger signal at the translation start site and one codon upstream of the stop site, and for single-nucleotide additions at the read start present in many libraries. Previous work used a similar conditional probability model for 5’-3’ bias in read coverage in RNA-seq based isoform deconvolution [22]. PRICE 2 groups reads into equivalence groups as described previously [21] and uses a Poisson generalized linear regression model to deconvolve the signal and estimate ORF activities under this generative model.

**Fig. 1.**
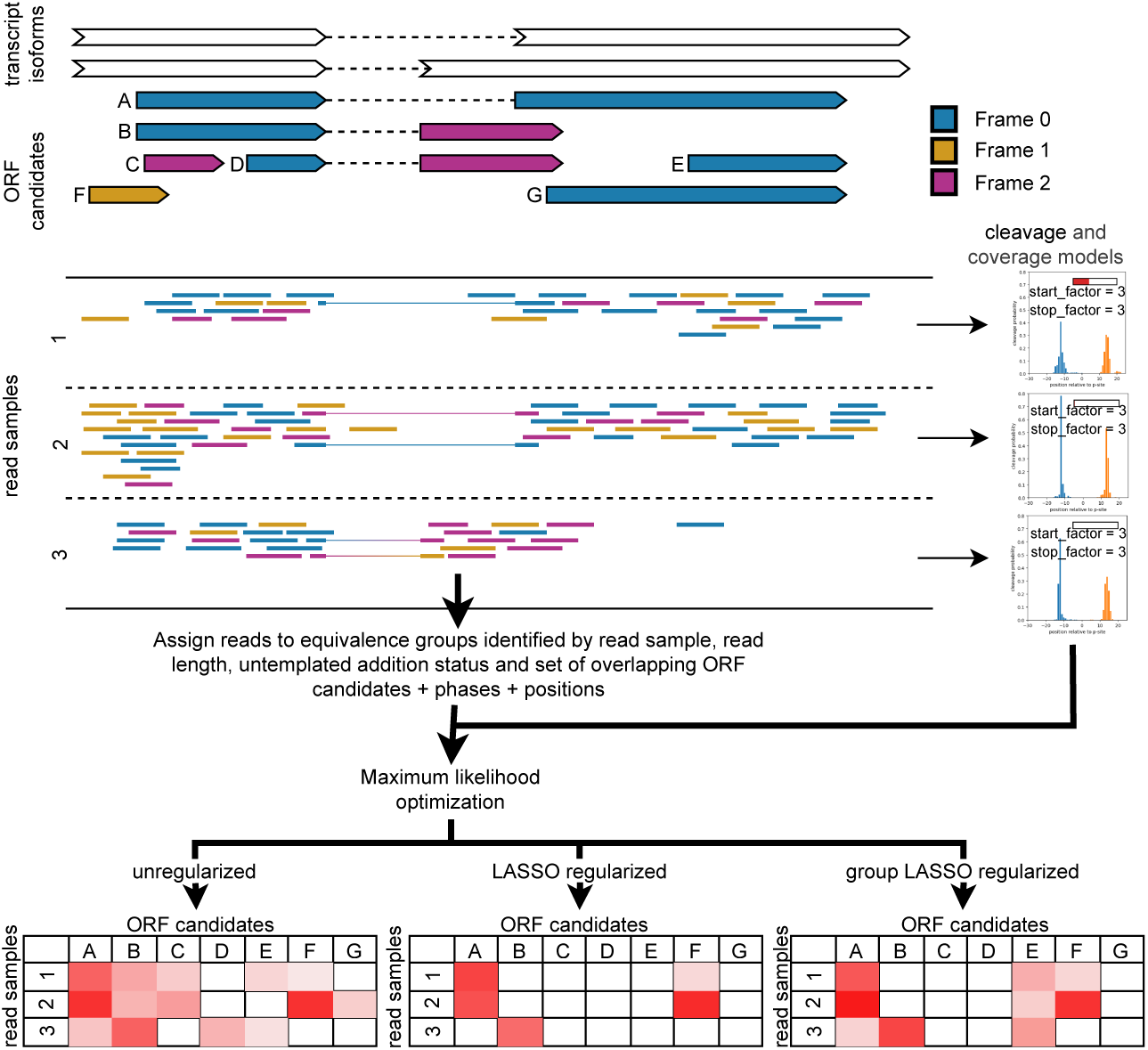
PRICE 2 applies a deconvolution algorithm based on a generative model for ribo-seq reads. Reads are assigned to equivalence groups and a group LASSO regularized robust Huber optimization is applied to estimate ORF activities over multiple ribo-seq samples. The group LASSO regularization induces sparsity in the solution and shares information between samples. The robust Huber approach minimizes the influence of outliers to reduce the number of false positives.

The conditional probabilities for the phase are computed from cleavage models introduced by PRICE 1. Cleavage models give the likelihood that a generated read has certain characteristics (read length, untemplated addition status and phase of the read start) [12]. Coverage models capture with scalar factors how much higher the read coverage is at translation start and stop sites. Analysis of many publicly available ribo-seq samples showed high variability in cleavage and coverage models (Supplementary Fig. 1). PRICE 2 therefore estimates these models separately for each sample (Fig. 2). This also allows PRICE 2 to be applied to ribo-seq data involving translation initiation inhibitors such as harringtonine or lactimidomycin, which by design have a very different coverage model with a much higher start factor (Supplementary Fig. 1 SRR13762831).

**Fig. 2.**
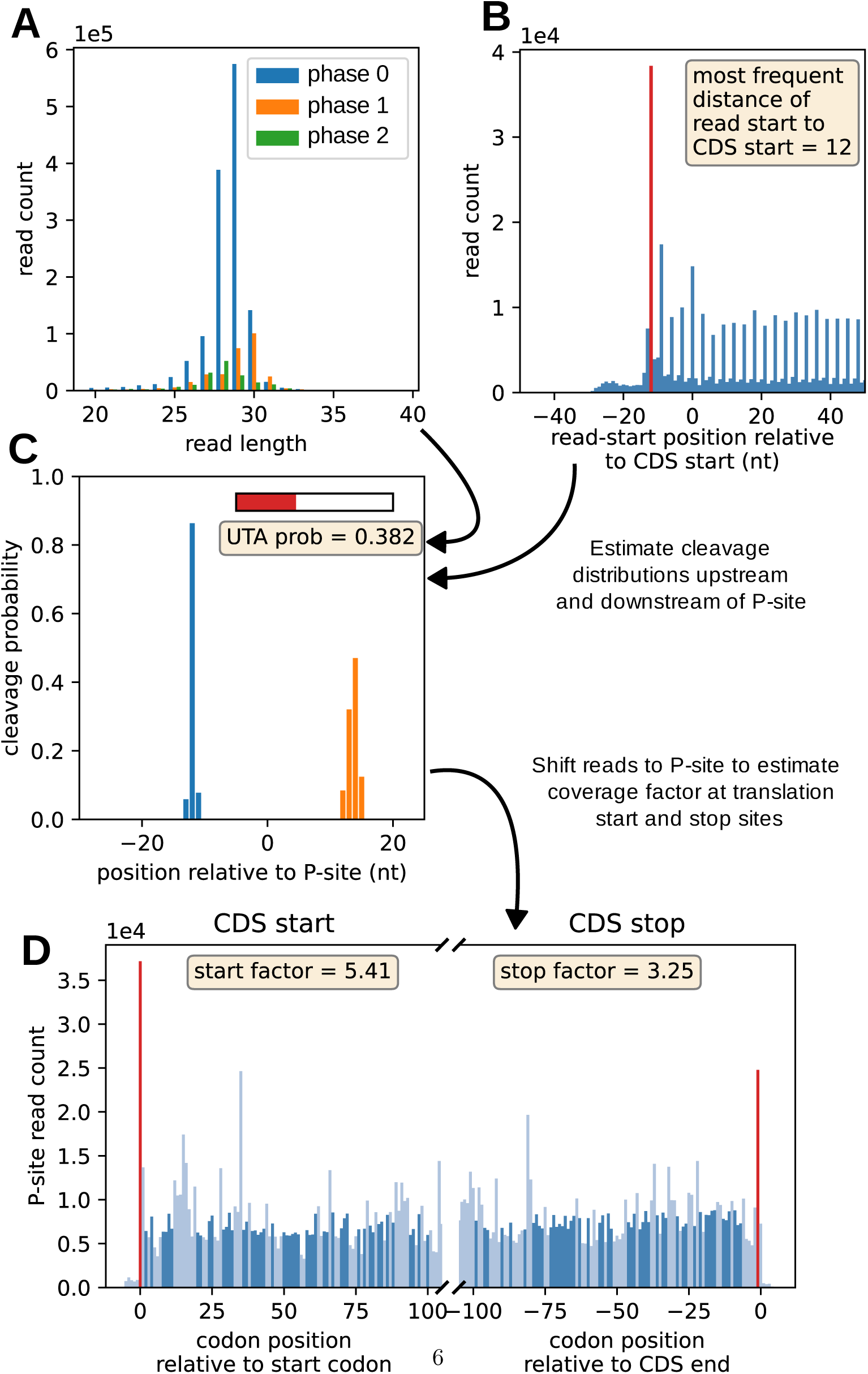
Characteristics of a ribo-seq sample are captured by cleavage and coverage models. The ribo-seq sample in this figure has the SRA id SRR12285191 **A:** Dataset wide statistics of read lengths and read starts to the underlying CDS phase. Only reads mapping to annotated protein-coding ORFs were used. **B:** Metagene plot showing cumulated read starts around translation initiation sites. **C:** Visualization of the cleavage model showing upstream (blue) and downstream (orange) cleavage probabilities from the ribosomal P-site and the probability of untemplated additions (UTA) at the 5’ end of the read. This model is estimated from data shown in **A** and **B** with an EM algorithm. **D:** Read P-site distributions around translation start and stop sites. Coverage at the start or directly upstream of the stop codon divided by 50% trimmed average of the other positions is used to compute the start or stop factor, respectively.

A major challenge for detecting weakly translated, short, and overlapping ORFs is the sparsity of the data. Previous work has addressed the low read coverage of individual ribo-seq runs by pooling multiple samples for ORF discovery [3, 16, 23–25]. Pooling overcomes sparsity but neglects useful information, e.g. that few reads observed in multiple samples provide more evidence for a translated ORF than the same number of reads in only one sample. We therefore apply a group LASSO regularization on the activity values [26], which makes full use of the data by sharing information across samples. In contrast to simple pooling, this ensures that low read evidence in each individual sample can still be detected when enough samples are available. A similar approach has been applied in the isoform deconvolution context [27]. The regularization also solves another major problem inherent to all such deconvolution approaches. The optimal solution typically involves many low but non-zero activity values caused by noise in the data. This makes it difficult to determine which ORFs are truly active and which are not. The group LASSO regularization overcomes this overfitting problem by inducing sparsity in the solution, i.e. by setting many activity values to zero.

Sequencing artefacts (such as amplification bias) can produce spurious read counts for some equivalence groups and therefore false-positive ORF calls. PRICE 2 handles this problem with a robust Huber Poisson model, which weights the influence of each equivalence group on the ORF activity. Equivalence groups that deviate more from the expected read count in intermediate optimization results receive smaller weights [28]. Short ribo-seq reads and the abundance of regions with high sequence similarity lead to many multi-mapping reads. With an EM algorithm, PRICE 2 distributes read weights over all locations according to the activities and fit of the ORFs over-lapping each mapping location. The M-step estimates ORF activities, and the E-step distributes the weight of each read over all locations it maps to. The M-step is computationally demanding even with few samples, because PRICE 2 solves a large Poisson regression problem with tens of thousands of data points and thousands of predictors. The number of equivalence groups/data points is high because of the large number of ORF candidates and because each read characteristic defines a separate equivalence group. PRICE 2 therefore runs the M-steps on each gene locus individually. Only the cheap E-step is computed unparallelized for all gene loci. The Poisson optimization is performed numerically. The online methods section gives a detailed description.

On simulated data, PRICE 2 detected active ORFs with high sensitivity even if they overlap another active ORF. Detection performance improved with more datasets and higher activity reaching accuracties of *>* 70% for iORFs translated at a level of 10% of the overlapping cORF (Supplementary Note 1, Supplementary Fig. 3).

To evaluate PRICE 2 on real data, we used a recent dataset of human B-cell lines with six standard ribo-seq samples, six QTI-seq samples and corresponding immunopeptidomic data [29]. We benchmarked PRICE 2 against most available state-of-the-art ribo-seq based ORF callers [12, 16–18, 24, 30–37] (Supplementary Table 1). Running bioinformatics tools is not always trivial because of bugs, incompatible dependency versions and insufficient or scattered documentation. More severely, benchmarks conducted by method developers are inherently prone to unconscious confirmation bias or over-optimism, as researchers may inadvertently invest more effort in fine-tuning their own tool while running competing tools with suboptimal defaults. To establish an unbiased and standardized evaluation, we leveraged recent advances in AI coding agents, which now complete tasks that take human experts more than one hour at an 80% success rate [38].

The AI agent ran all tools successfully. Not all ran on the first attempt because of misspecified dependencies or bugs. We instructed the agent to fix such issues without human intervention and to report all steps. PRICE 2 was the fastest tool at 38 minutes (Supplementary Table 1). ribotricer produced the most predicted ORFs (n=236,831), followed by PRICE 2 hrt (n=135,440) (Fig. 3A). ORF-RATER produced the fewest (n=14,088). PRICE 2 produced more ORFs than most other tools (n=109,247). It detected more +2 uoORFs (n=2,718) and +2 iORFs (n=8,850) than any other tool (for ORF type definitions see Supplementary Fig. 2). Only RiboBA detected more uORFs (n riboba=20,303, n price2=13,463) and +1 uoORFs (n riboba=3,206, n price2=2,719). Only ribotricer produced more +1 iORFs than PRICE 2 (n ribotricer=31,758, n price2=20,151). PRICE 2 detected significantly more +1 iORFs (n=20,151) than +2 iORFs (n=8,850). Adding the harringtonine (hrt) treated samples for PRICE 2 raised the number of detected ORFs, especially for uORFs (n price2=13,463, n price2 hrt=21,415) and uoORFs (n price2=5,437, n price2 hrt=10,702). These numbers are no measure of prediction quality because the ORF sets mix true and false positives.

**Fig. 3.**
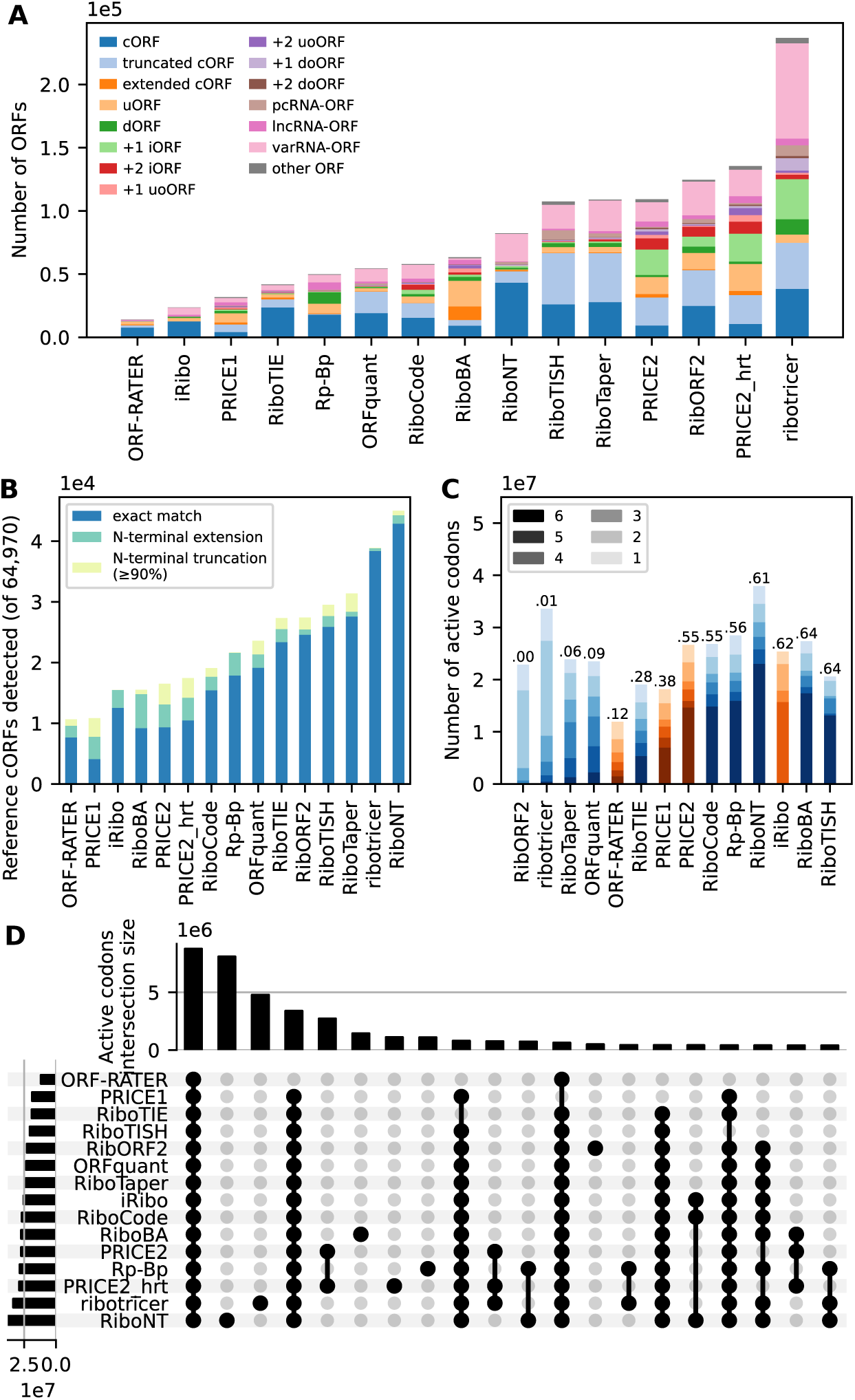
Evaluation on real data compared to other tools. **A**: Number of detected ORFs by type for tools applied on the same samples. **B**: Number of cORFs from the reference annotation that was detected or approximately detected. **C**: Reproducibility of active codon calls across the 6 ribo-seq samples. Darker shades indicate detection in more samples. Numbers on top of the bars are the multi set Jaccard index. Tools that natively operate on multiple samples (highlighted in red) were applied on each sample individually for a fair comparison. RibORF2 was only applied on 5 datasets and iRibo on 4 because they filter samples on a periodicity threshold. **D**: Set comparison of active codons. Only the 20 largest intersections are displayed.

We evaluated how many of the 64,970 cORFs from the reference annotation each tool rediscovered exactly or approximately (Fig. 3B). We defined approximate rediscovery as an N-terminal extended cORF or an N-terminal truncated cORF with at least 90% of the cORF length. ORF-RATER predicted the fewest cORFs (n=10,637) and RiboNT the most (n=45,004). PRICE 1 and PRICE 2 placed at the lower end with n=10,838 and n=16,496 respectively. Recovery of cORFs is a poor measure of prediction quality because many tools implement statistical or machine learning models that were trained on or adjusted with cORFs as a positive set, even though many cORFs in these samples will not have been active. The tools then detect ORFs resembling their training set, which yields false positive cORF detections. The problem is compounded because many gene loci have several cORFs annotated that overlap in large parts, which gives a strong frame signal for many cORFs even when only one is active.

High reproducibility of active codons across similar samples is necessary, though not sufficient, for accurate ORF detection (Fig. 3C). For a fair comparison we ran tools with native multi-sample support on each sample individually. RibORF2 and iRibo did not run on all 6 samples because of periodicity thresholds they apply. On this metric RiboTISH and RiboBA performed best among the tested tools, with a multiset Jaccard index of 0.64 each, followed by iRibo at 0.62 and RiboNT at 0.61. RiboNT uses the codon distribution in addition to the RPFs, which explains its high number of cORFs and its high reproducibility. RibORF2, ribotricer and RiboTaper showed very low reproducibility. PRICE 2 had a multiset Jaccard index of 0.55.

Comparing tools at the ORF level is complicated because they often predict different variants of the same ORF, e.g. with different start codons. Comparing active codons (codons that are part of at least one predicted ORF) shows the agreement between tools more clearly (Fig. 3D). While ribotricer produced the largest set of active ORFs, RiboNT produced the largest set of active codons (n=37,927,216), followed by ribotricer (n=33,584,346) and PRICE 2 hrt (n=28,861,101). Rp-Bp produced a large set of active codons (n=28,479,045) but a rather small set of active ORFs (n=49,872). ORF-RATER produced the smallest set of active codons (n=11,867,176), as it did for active ORFs. PRICE 2 called n=27,356,667 codons active. The largest intersection was the set called active by all tools (n=8,768,089), showing high agreement. A similarly large set was called active only by RiboNT (n=8,090,811), and another large set only by ribotricer (n=4,780,082). ORF-RATER missed many codons that all other tools called active (n=3,388,118), and a slightly smaller set was called active only by PRICE 2 and PRICE 2 hrt (n=2,722,045). Importantly, the tools that detect relatively few cORFs missed no large set of codons, which suggests that they usually missed only cORF isoforms rather than all cORFs of a gene locus.

Visual inspection of the ORF predictions on individual loci confirmed or explained many of the systematic-level observations (Supplementary Fig. 5). PRICE 2 detected many short ORFs and often predicted multiple active start codons for a single stop codon, though fewer than tools such as RiboNT or ribotricer. Agreement between PRICE 2 with and without harringtonine-treated samples was high, and both versions detected iORFs with strong read evidence that the other tools missed.

Without a ground truth for the real data, none of these comparisons directly show that PRICE 2 detects ORFs more accurately than the other tools. To assess accuracy we leveraged immunopeptidomics data from the same cell lines [29]. We applied Peptide-PRISM [9], which employs de-novo peptide sequencing [39] to identify presented peptides in a database agnostic manner. Peptide-PRISM then searches the identified sequences in the entire six-frame translated human genome and filters them for a desired false discovery rate (FDR) via a stratified target-decoy approach. This controls the FDR within each predefined category, but gives weak sensitivity for large strata such as all intronic or intergenic sequences. Sensitive Peptide-PRISM results therefore depend on strata that contain many presented peptides relative to their size. Strata were defined in the following order: the predicted ORF products for a given tool first, then the canonical proteome, and finally the default Peptide-PRISM databases such as 3-frame translation of the 5’ untranslated region, the 2 out-of-frame translations of coding regions, 3’ untranslated regions and non-coding RNAs. We ran Peptide-PRISM for each tool at an FDR of 10%. We assigned peptides to ORF types hierarchically, e.g. a peptide that could stem from an active cORF was never assigned to a truncated cORF even when it overlaps. The number of peptides detected in the tool-predicted ORFs was similar for most tools (Fig. 4A). The fewest fell in the ORFRATER ORFs (n=3,396) and the most in the RiboNT ORFs (n=4,163). Most peptides could be assigned to cORFs, except in PRICE 1, where a large fraction was assigned to extended or truncated cORFs. Even in the PRICE 2 variants, which detected fewer cORFs than most other tools, most peptides were assigned to cORFs (77%). This shows that many of the active cORFs in the other tools may be false positives. To analyse cryptic peptides we reran Peptide-PRISM with another order of the strata: the canonical proteome first, then the tool predicted ORF products, and finally the default Peptide-PRISM databases (Fig. 4B). The number of cryptic peptides detected in tool ORFs varied widely, from 13 for iRibo to 101 for PRICE 2 hrt. Even without harringtonine-treated samples PRICE 2 outperformed the other tools (n=91). The next best was ribotricer (n=71), which predicted by far the most ORFs of any tool.

**Fig. 4.**
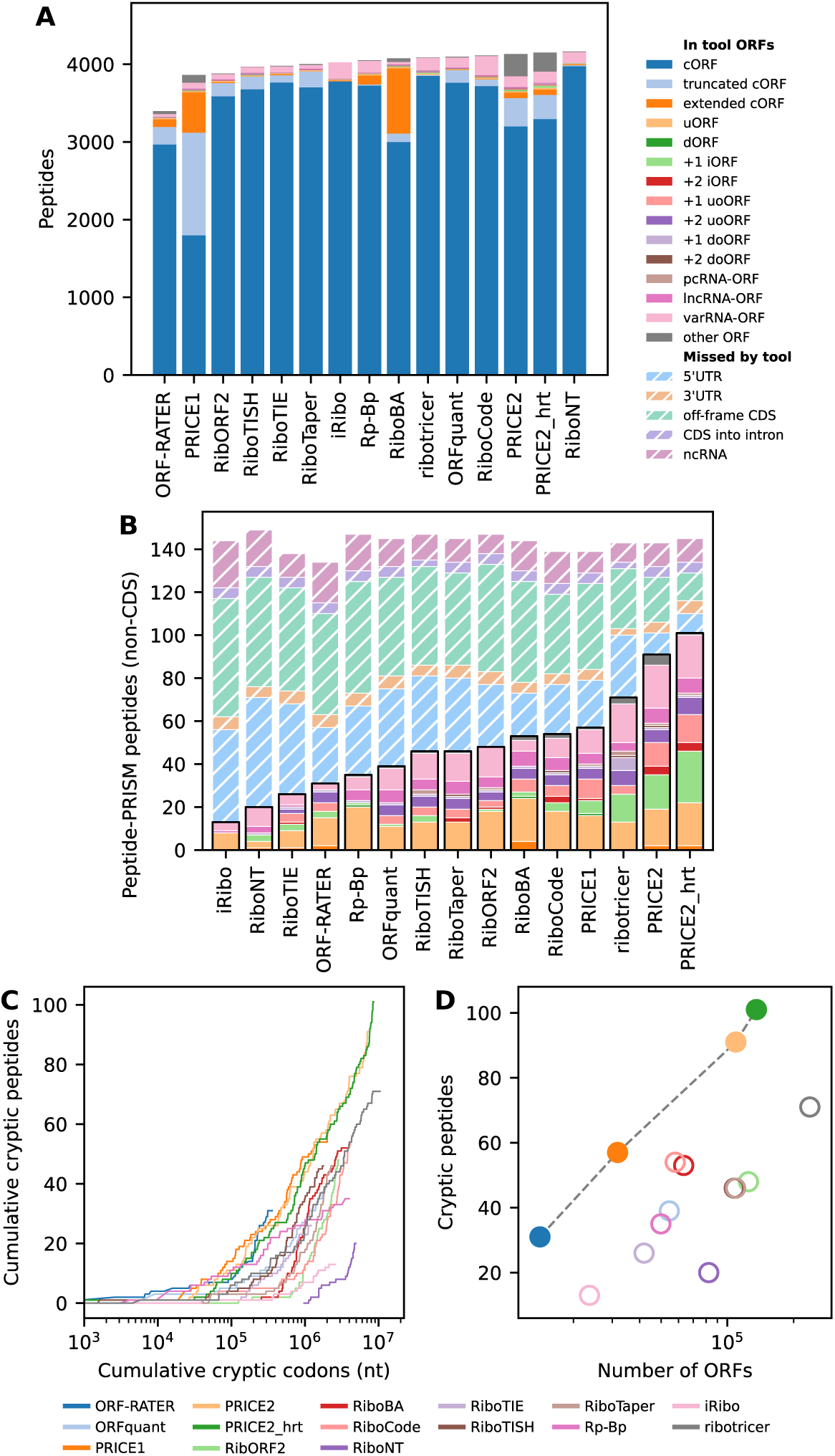
Evaluation with immunopeptidomics as a complementary method. **A**: number and type of peptides that could be found in the proteins from each tools. Peptides were assigned to one ORF type in a hierarchical manner. **B**: number and type of cryptic peptides that could be found in the from each tool and in other strata by Peptide-PRISM. **C**: Cumulative explained peptides vs codon count. ORFs from each tool were sorted by an average activity or reliability score starting with the most active/certain. **D**: A Pareto plot for the tools and databases where a high number of cryptic peptides and a low number of called ORFs are the values to be optimized. The Pareto front is highlighted. The RiboTISH marker is covered by the RiboTaper marker.

The PRICE 2 ORFs explained more peptides than all other tools for iORFs, uoORFs and varRNA-ORFs, whereas RiboBA and Rp-Bp explained slightly more uORF peptides (n=20 each vs. n=17 for PRICE 2). The largest difference was in +1 iORFs, where PRICE 2 hrt detected 24 peptides and the next best tool ribotricer only 13. Interestingly, only 4 peptides were detected in +2 iORFs for both PRICE 2 and PRICE 2 hrt.

Only a small fraction of truly translated ORFs is expected to yield a peptide in immunopeptidomics data, since many translated proteins are not presented at the cell surface and many peptides are not detectable by mass spectrometry. Peptide presentation is also not necessarily proportional to the translation level of the corresponding ORF, and many presented peptides may well originate from ORFs translated at levels too low to give any detectable signal in ribo-seq data. Comparison with immunopeptidomics data therefore cannot estimate the absolute performance of a ribo-seq tool. It does, however, allow a comparison of the relative sensitivities of the different tools at a fixed FDR. In summary, our results show that PRICE 2 is more sensitive than the other tools for the detection of ncORFs, and not merely because it predicts more ORFs. It is in particular more sensitive than all other tools for the most challenging cases (iORFs).

We hypothesized that a higher activity cutoff for predicted ORFs could yield an even greater number of detected peptides, because a smaller sequence database gives Peptide-PRISM greater statistical power. To test this, we plotted the cumulative number of identified peptides against the cumulative number of codons for all activity threshold values for PRICE 2. For the other tools we used activity values where available and significance or certainty scores otherwise. Codons at all activity levels contribute to the detected peptides, and no activity threshold would yield a similar number of detected peptides with fewer codons (Fig. 4C). The curve is nonetheless steeper for more active codons, which therefore contribute more to the peptide count. ORF-RATER, PRICE 1 and PRICE 2 show the steepest incline and thus the highest cryptic peptide to cryptic codon ratios. We propose to treat ORF calling from ribo-seq data as a Pareto optimization problem, maximizing the number of cryptic peptides detected from immunopeptidomics data while minimizing the number of ORFs. ORFRATER, PRICE 1 and the PRICE 2 variants form Pareto-optimal solutions regarding these quantities (Fig. 4D).

We evaluated a subset of the tools with manual tool invocations giving results comparable to the agentic workflow (Supplementary Note 2, Supplementary Fig. 4). The nuORFdb [40] is a comprehensive database of ncORFs, built by hierarchically applying RibORF [24] and PRICE 1 to 29 ribo-seq samples from a range of tissues and cell lines. One of its main uses is scanning immunopeptidomics data for cryptic peptides, and the better performance of PRICE 2 in this application suggests that a database built from its predictions would outperform the nuORFdb. We therefore assembled a diverse, high-quality set of public ribo-seq samples. We queried the Sequence Read Archive (SRA) and European Nucleotide Archive (ENA), filtered the hits, then downloaded and mapped the 3,113 high-quality samples that remained (Fig. 5A). Clustering by gene counts gave 45 clusters corresponding to tissue or cell types (Fig. 5B), and we applied PRICE 2 to the four best samples of each cluster, chosen by read count and periodicity (Fig. 5C). To see how much the database depends on the number of input samples, we also built *ORF-stock-45* from the single best sample of each cluster. We evaluated the *ORF-stock*, *ORF-stock-45*, nuORFdb and TransCODE databases on the immunopeptidomics data used for the tool comparison. The TransCODE databases were built for specificity. Their authors merged ncORFs from nine studies and filtered them with ribo-seq-based translation-evidence scores (“P-sites in frame”, “Uniformity” and “Dropoff” at translation start and stop sites), yielding 10,127 (primary) and 28,359 (comprehensive) ORFs [41].

**Fig. 5.**
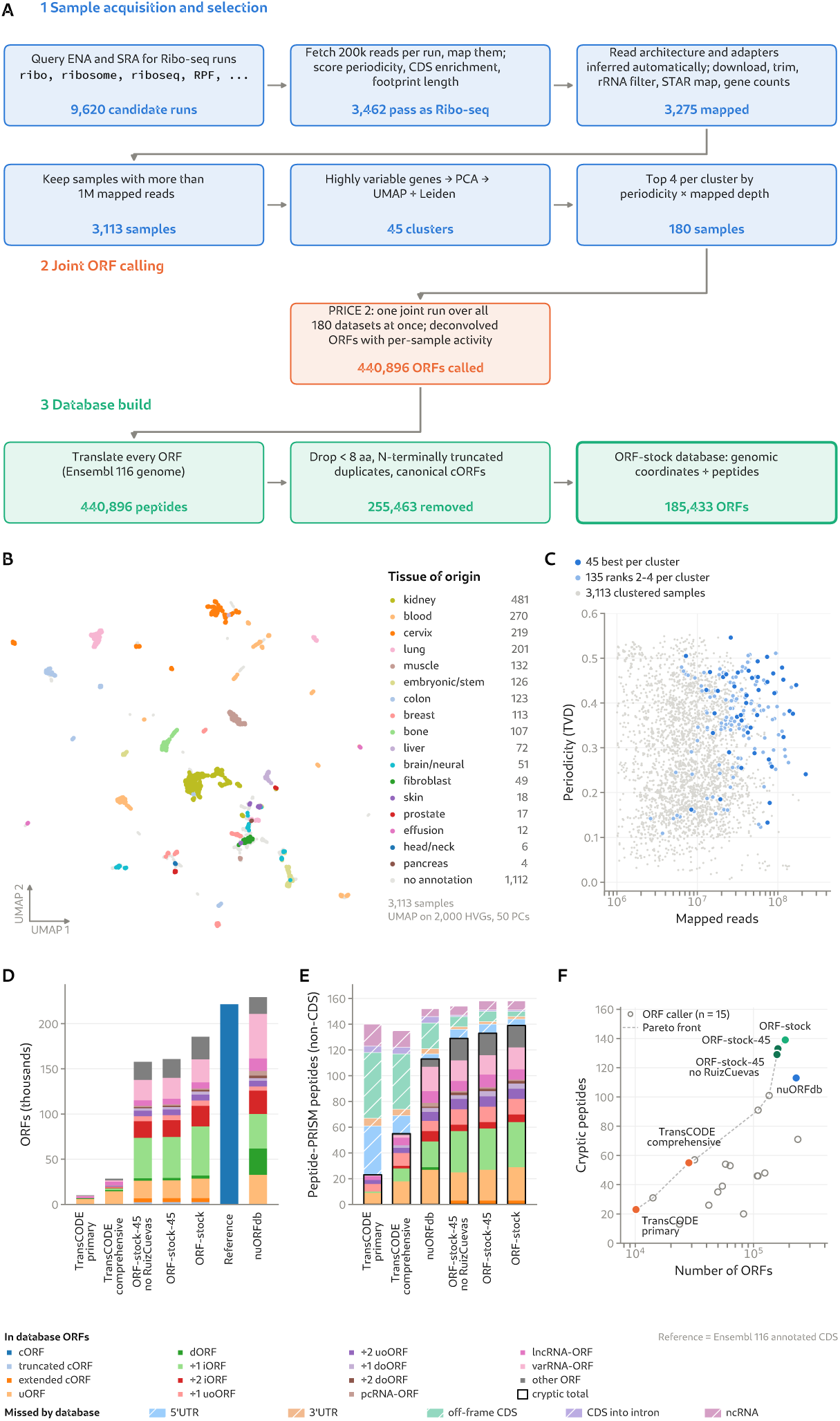
The *ORF-stock* database was created by applying PRICE 2 to a diverse high-quality set of ribo-seq samples. **A**: Workflow to retrieve and characterize ribo-seq samples from public repositories **B**: UMAP of ribo-seq samples based on read counts. Colours represent tissue type as inferred from the meta data **C**: Scatter plot with Read start periodicity vs number of mapped reads for each ribo-seq sample. Samples used for *ORF-stock* (best four per cluster, n=180) are highlighted; dark: best sample per cluster (n=45), used for *ORF-stock-45*. **D**: Number of ORFs in the databases by type; Reference: coding ORFs annotated in Ensembl 116 **E**: Number of cryptic peptides detected by Peptide-PRISM in the immunopeptidomics data from [29] for each database, coloured by ORF type. Hatched segments above the frame show non-CDS Peptide-PRISM peptides not explained by a database ORF, by Peptide-PRISM category **F**: Pareto plot with many cryptic peptides and few ORFs being better solutions. Databases are displayed in colour and labelled; *ORF-stock-45* : built from the best sample per cluster only, *ORF-stock-45 no RuizCuevas*: as *ORF-stock-45* with the ribo-seq sample matching the immunopeptidomics data replaced. ORF callers displayed in grey.

All four *ORF-stock* samples of one cluster came from the same study as the immunopeptidomics data [29], and the best of them was also used for *ORF-stock-45*. To test whether performance depends on these matched samples, we generated a version of *ORF-stock-45* in which this sample was replaced by the best sample of its cluster from a different study (*ORF-stock-45 no RuizCuevas*).

*ORF-stock* contains fewer ORFs than nuORFdb (n=185,433 vs. n=229,249), with more +1 iORFs but fewer varRNA-ORFs, dORFs and lncRNA-ORFs. Using four times as many samples increased the database size only moderately (*ORF-stock-45* : n=160,743). The TransCODE databases are far smaller, matching their design goal of specificity over sensitivity (Fig. 5D). Peptide-PRISM detected more cryptic peptides in *ORF-stock* (n=139) than in *ORF-stock-45* (n=133), nuORFdb (n=113) and the TransCODE databases (primary: n=23; comprehensive: n=55) (Fig. 5E). Replacing the matched sample cost *ORF-stock-45* only four cryptic peptides (*ORF-stock-45 no RuizCuevas*: n=129), so its advantage over nuORFdb does not depend on ribo-seq data from the cells analysed by immunopeptidomics. All databases except nuORFdb are Pareto-optimal, even when the ORF calling tool results are included (Fig. 5F).

## Discussion

Ribo-seq has enabled great strides in detecting translated ORFs in the human and other genomes. Most importantly, it revealed that previous annotation efforts based on evolutionary conservation, sequence information and mass spectrometric evidence missed many short ORFs. However, de-novo sequencing of MHC-I derived peptides has shown that all existing tools, including our PRICE 1, lack sensitivity. Their approaches inherently struggle to detect short and lowly translated or cORF-overlapping frameshifted ORFs.

Here we introduce PRICE 2, which overcomes this limitation with a generative model well established for RNA-seq-based isoform deconvolution and used by state-of-the-art tools such as Salmon [42]. PRICE 2 can process multiple ribo-seq samples at once, estimating ORF activities per sample while a group LASSO penalty lets the samples inform each other. Cleavage and coverage models capture the characteristics of each sample, so PRICE 2 uses the relevant signal optimally. It also extends seamlessly to QTI-seq data, which uses antibiotics to stall ribosomes at initiation sites, effectively enriching reads at these positions.

Benchmarking computational analysis tools for ribo-seq data is difficult for two major reasons: First, comparing to previous tools is highly challenging because of outdated tools or dependencies and insufficient documentation. We show how language model based coding agents can ease this task while reducing bias in the comparisons. Second, evaluating the absolute and exact sensitivity and specificity of any tool is virtually impossible because no ground-truth dataset exists. The binary classification of ORFs as active or inactive may also be a strong simplification, since noise in the translation process may cause many ORFs to be translated only occasionally. We used immunopeptidomics data as an orthogonal method to evaluate the ORF calls of the different tools. PRICE 2 ORFs explain a larger fraction of peptides than those of the other tools, demonstrating higher sensitivity. In particular, peptides corresponding to iORFs were almost absent from the other tools’ predictions. These are among the most challenging cases, as iORFs are typically short and overlap the typically much stronger translated cORF. Of note, peptide identification strictly controlled the FDR for all tools. Thus, the superior sensitivity of PRICE 2 does not come at the cost of sacrificed specificity.

Due to technical (adapter ligation and amplification bias, read mappability) and biological (varying translation speed, ribosome pausing) factors, ribo-seq read coverage within an active ORF is not uniform, which can cause false positive or false negative ORF calls. PRICE 1 assumed that RNase cleavages, and therefore read starts and lengths, follow the same distribution for each active codon. PRICE 2 uses this model differently and makes a much weaker assumption, namely that cleavages should be consistent with the distribution across an entire equivalence group. It also applies a robust Huber Poisson regression to further reduce the impact of spurious signals. In principle, PRICE 2 could model further signals in ribo-seq samples, e.g. translation velocity (leading to different codon occupancies) or amplification bias. The model could then draw information from reads at more nucleotide positions instead of discarding it for not fitting. Unique molecular identifiers can avoid sequencing biases, and predictive models to estimate ribosome dynamics already exist [43].

The “dark proteome” has drawn a lot of attention as an untapped source of tumor-specific antigens [11], prompting major international efforts such as the Cancer Grand Challenges “Dark Proteome” initiative supported by Cancer Research UK and the National Cancer Institute [44]. State-of-the-art immunopeptidomics relies on *de novo* sequencing with reference-guided searches to identify presented cryptic peptides [9, 29, 39]. As massive sequence search spaces such as the full, three-frame translated human transcriptome severely impair statistical power, a high-quality ORF sequence database is essential. nuORFdb [40], which was built from 29 samples using RibORF [24] and an unfiltered version of PRICE 1, has been widely adopted for that purpose. By applying the improved deconvolution and regularization of PRICE 2 to a diverse compendium of 180 quality-filtered datasets, we established *ORF-stock*. ORFs in *ORF-stock* were significantly enriched in tumor-associated peptides and markedly outperformed nuORFdb, especially for the most challenging cases (iORFs), providing a comprehensive resource for cryptic antigen discovery.

## Online methods

### Cleavage models

Ribo-seq samples have a characteristic read length distribution and each read length has different propensities to start in certain phases (0/1/2, position relative to a codon). These characteristics emerge from the 3 nt translocations of the ribosome during translation and from cleavage probabilities of the RNase upstream and downstream of the ribosomal P-site. Furthermore, a fraction of ribo-seq reads have untemplated additions (reverse transcriptase nucleotides) at the 5’ end: single nucleotides present that are neither part of the biological sequence nor constant length technical sequence. Cleavage models can be estimated from a set of characterized reads and the coverage profiles proximal to translation start sites using an EM algorithm. They can be used to compute the likelihood of a given read characteristic (read length, untemplated addition status, and phase of the read start) [12]. Cleavage model estimation for PRICE 2 is identical to PRICE 1. Briefly, each sample was randomly downsampled to at most 10 million mapped reads, and all reads of that subsample that map uniquely within an annotated CDS were used to fit the model. The reads were characterized for read length, untemplated addition status, and reading frame relative to the overlapping coding sequence from a reference annotation. Reads that were not uniquely mapped, mapped to multiple coding sequence regions coding in different frames, or mapped to no coding sequence were discarded.

### Coverage models

In many ribo-seq samples, coverage is elevated at positions where the P-site of the ribosome is at an ORF start site and one codon upstream of the stop site. The coverage model is defined by a start and a stop factor indicating how much higher the coverage is relative to the average coverage at positions within ORFs. Coverage models were estimated using the same downsampled BAM. Only reads that could be assigned to one in-frame P-site with a probability *>* 0.8 according to the cleavage model were considered. The start factor was computed as the number of reads mapping to the start codon divided by the 50% trimmed average number of reads mapping to the first 300 positions (excluding the start). The equivalent approach was applied for the stop factor. We note that coverage models naturally extend to translation initiation sequencing data (QTI-seq).

### ORF deconvolution

Many tools that detect ORFs from ribo-seq data generate a set of ORF candidates and compute characteristics such as 3 nt coverage periodicity or coverage uniformity to find translated ORFs. One caveat of this approach is that the signal for overlapping ORFs is convoluted, leading to poor detection of iORFs. We introduce a statistical model to address this limitation, inspired by isoform deconvolution tools for RNA-seq data.

#### A generative model for ribo-seq data and the maximum likelihood estimator

For an ORF *j* consisting of *n* codons (excluding stop), let *R_j,i_* be the number of reads generated by the ribosomes with the P-site at codon position *i* while translating ORF *j*. Let *γ_i_* be a specific scalar factor for each position. Here we set *γ*_1_ and *γ_n_*to the start and stop factor, respectively, and *γ_i_* = 1 for all other codons. We assume that *R_j,i_* has an identical expectation *β_j_*, called activity of ORF *j*, for all internal codon positions *i* 2*, . . . , n* 1 , which is scaled by the start or stop factor, respectively, for the first and last position:

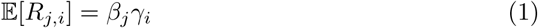

Accordingly, the expected number of reads for a subset of codon positions from ORF *j* is

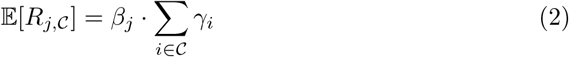

Let *R_j,l,f,u_* be the number of reads with length *l*, phase *f* , and untemplated addition status *u* generated due to translation from any position in ORF *j*, and *p*(*l, f, u*) be the cleavage probability for length *l*, phase *f* , and untemplated addition status *u*. Then the expectation of *R_j,l,f,u_*is

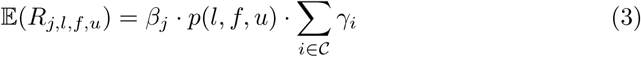

While most reads originate from a ribosome that translated a particular ORF *j*, for many reads there are several candidate ORFs that could have generated the read (mainly because of overlaps among the ORFs). The same is also true for the isoform deconvolution problem of RNA-seq, where reads are assigned to equivalence groups that are defined by all isoforms that they are compatible with [21]. For ORF deconvolution, computing likelihoods also depends on the cleavage and coverage models. Thus, an equivalence group *G ∈ G* is defined by a read length *l* and an untemplated addition status *u*, and a set of codon positions *_j_* from one or several ORFs *j J_G_*. These ORFs belonging to *G* can overlap with identical phase relative to each other or with distinct phases. Thus, each ORF has a phase *f_j_* relative to the equivalence group. There are also separate equivalence groups according to coverage positions (start, one upstream of stop, or other). The expected read count for an equivalence group can be computed as the sum of the expected read counts for the overlapping ORFs. Importantly, if the ORFs code in different frames, different frame parameters must be applied for the cleavage model.

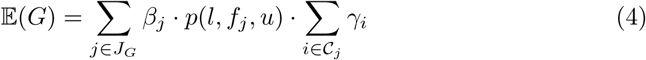

We assume that the read count of an equivalence group is Poisson distributed. Denoting the observed and expected read counts of equivalence group *G* as *r_G_* and *δ_G_* = E(*G*) respectively, we can compute a likelihood for one equivalence group:

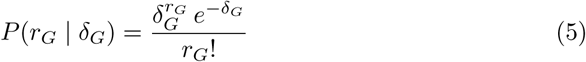

Considering all equivalence groups for one ribo-seq sample, we can compute the overall likelihood:

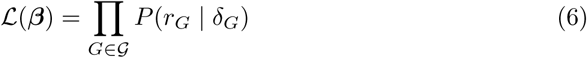

For multiple ribo-seq samples *d D*, we must account for the fact that equivalence groups can differ between samples due to different cleavage models:

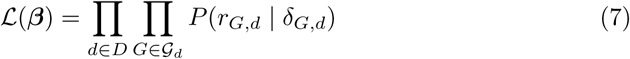

The optimal activities maximize this likelihood. In practice, we minimize the negative logarithm of this likelihood. The read count factorial term does not affect the location of the minimum and can therefore be discarded:

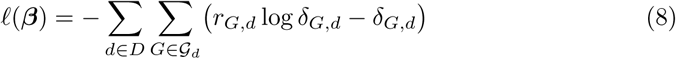

This optimization function is convex [21] and can therefore be solved by any convex optimization algorithm.

#### Group LASSO Regularization

Since we expect only a small fraction of all ORF candidates *J* to be active, we regularize the objective function with a group LASSO penalty [26]. Each group contains the activity values *β* belonging to one ORF across all sequencing runs. The group LASSO not only induces sparsity in the solution but also shares information between sequencing runs, meaning that a lowly active ORF that would be regularized to zero if optimized for each run individually may be estimated as active across all runs jointly.

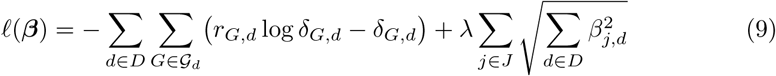

#### Poisson regression

The model described in the previous sections is equivalent to a Poisson regression model with an identity link function and no intercept. In this view, each equivalence group corresponds to a sample and each (ORF, sample) activity *β_j,d_* to one feature. Accordingly, the design matrix *X* has one row per equivalence group *G* and one column

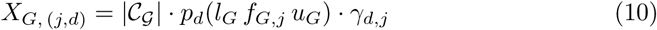

where *|C_G_|* is the number of positions in *G*, *p_d_* is the cleavage model of sample *d*, *l_G_*, *f_G,j_*, *u_G_* are the read length, phase relative to ORF *j*, and untemplated addition status of group *G*, and *γ_d,j_* is the coverage factor for ORF *j*.

Since most ORFs do not overlap most equivalence groups and each equivalence group is only valid for one sequencing run, this matrix is very sparse. PRICE 2 stores the matrix as a scipy sparse matrix and the estimated activities and equivalence group read counts as numpy arrays, enabling highly efficient computation of the likelihood and loss.

#### Robust Huber regression

PRICE 2 applies a robust Huber regression to mitigate the influence of outliers [28]. In Huber regression, the optimization function is first fit with unweighted data points. From the initial fit, the Pearson residuals for each data point are computed; data points with large residuals are down-weighted and the next round of minimization is run. This process is repeated until the relative change in all weights falls below a threshold. The Pearson residual and Huber weight function are:

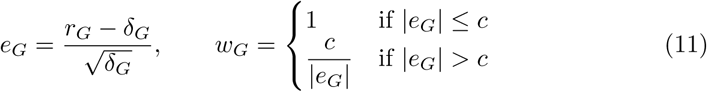

where *c* is the robustness tuning constant.

#### Multi-mapping read distribution

Running the Poisson regression on all genes as one optimization is computationally not tractable. Instead PRICE 2 runs it on each gene locus separately. This introduces the problem that reads that map to multiple genomic locations need to be considered in multiple disjoint optimizations. PRICE 2 distributes these multi-mapping reads with an outer EM loop. The previously described Poisson optimization just estimates ORF activities and is the M-step. In the E-step the multi-mapping reads are fractionally distributed to the mapping locations with weights computed according to the activity and fit of the overlapping ORFs. This EM loop is stopped when the maximum absolute activity change over the ORFs is below a threshold.

### The Workflow to detect ORFs

#### Computing characteristics of ribo-seq runs

A subset of 10 million mapped reads is subsampled from each sample to compute cleavage and coverage models, using a reference annotation. The phase of the read start relative to the overlapping coding sequence (CDS) and the distance of the read start to the overlapping CDS start and stop sites are computed. Only reads that uniquely map to one CDS frame are considered. Cleavage models are computed from the characterized reads and the coverage profile of read starts around CDS start and stop sites, as described in [12]. The number of possible read lengths that can be generated from a cleavage model is proportional to the number of equivalence groups. To reduce the number of equivalence groups, we regularize the cleavage models by retaining only the highest upstream and downstream probabilities such that they sum to at least 0.9, and rescale the retained cleavage distances to sum to 1.

Coverage models consider only a start and a stop coverage factor. These factors are computed by estimating the P-site position relative to the start and stop of the overlapping CDS for the 100,000 sampled reads using the cleavage model. The start factor is computed as the P-site coverage over the start codon divided by the 50% trimmed average P-site coverage over the first 100 codon positions (excluding the start). The stop factor is computed equivalently as the P-site coverage over the position one codon upstream of the stop divided by the average of the last 100 codon positions. If the computed start or stop factor is smaller than one, it is set to one.

#### Generating gene loci

Gene loci are generated according to the reference annotation. Annotated transcript intervals are stored in an interval tree. Clusters of overlapping transcripts and transcripts nested within introns of other transcripts form a locus.

#### Collecting read information for efficient processing

The deconvolution step in PRICE 2 is parallelized over gene loci but uses read information from all samples for each locus. To efficiently access the necessary data for each locus, PRICE 2 implements a data collection step that runs in parallel over the samples. In this step, equivalent mappings are grouped and the necessary information for each mapping is determined: genomic mapping intervals, observed untemplated additions, and mapping count (from the grouping). This data is binarized, compressed, and saved in a SQLite3 database with one entry per gene locus per sample.

#### Isoform filtering and generating ORF candidates

For each gene locus, all annotated isoforms are considered. The isoforms are then filtered to retain a small set that explains most of the observed reads across all samples. We apply a greedy algorithm that selects the isoform explaining the most reads, adds it to the retained set, and removes it from the candidate set. This process is repeated on reads not yet explained by any retained isoform until the number of explained reads falls below a threshold for all remaining isoforms. Candidate ORFs are generated from the sequences of the retained isoforms. All combinations of start codons (default: ATG, CTG, GTG, TTG) and stop codons (default: TAA, TAG, TGA) forming an ORF (i.e., no inframe stop codons between them) are considered ORF candidates. ORF candidates are mapped back to genomic coordinates and identical ones (from different isoforms) are collapsed.

#### Generating equivalence groups

PRICE 2 generates equivalence groups defined by the sequencing run, read characteristics (length, untemplated addition status), and the overlapping ORFs with the corresponding phase and coverage position. Equivalence groups have a length (i.e., the number of nucleotide positions at which a read of that equivalence group could be generated). When generating equivalence groups and assigning reads, PRICE 2 accounts for the fact that reads spanning splice junctions can only be generated by ORFs that also span those junctions. To generate equivalence groups, PRICE 2 constructs a splice graph from all ORF candidates. The genomic positions corresponding to edges in the graph are shifted by likely cleavage distances. Equivalence groups are generated from each node (or the length of an existing equivalence group is incremented) by considering all overlapping ORFs and all possible read characteristics. PRICE 2 also accounts for the fact that P-site positions close to a transcript start or end cannot generate reads, as the resulting reads would be too short and would be filtered.

#### Assigning reads to equivalence groups

Mapped reads are assigned to equivalence groups by determining the read characteristics and the overlapping ORFs with the corresponding phase and coverage position. These characteristics serve as the key for the dictionary of equivalence groups. When a read is assigned to an equivalence group, the read count is incremented by the number of reads in the group of equivalent mappings. When a read overlaps the start or stop site of an ORF, it is possible that only an unlikely P-site position within the read fits the ORF. Whether such a read is assigned to an ORF is determined as follows: the likelihood that the read fits the ORF assuming complete overlap is computed, and the partial-overlap likelihood is computed. If the ratio of the partial-overlap likelihood to the complete-overlap likelihood exceeds a threshold, the read is assigned to the ORF.

#### Heuristic ORF filters

The loss function presented in the previous section captures many aspects of ribo-seq samples, but it has the drawback of being computationally demanding to optimize due to the large number of equivalence groups. Furthermore, we have observed that it retains many false positives even under strong regularization. To make the estimation tractable and to improve PRICE 2’s accuracy, we apply a number of filters on predicted ORFs before and after the deconvolution step.

#### Coverage filter

We applied a coverage filter that only considers reads that can be assigned to one frame with high confidence. We computed the average coverage of these reads for each ORF candidate and each sample. We discarded all ORF candidates where the maximum over all samples is below a threshold.

#### Deconvolution filter

We applied a computationally efficient deconvolution filter on smaller subsets of ORF candidates before running the full deconvolution. These subsets consist of ORF candidates that share the same stop codon and have compatible exon-exon junctions. Applying deconvolution to these subsets allows considering only well-fitting reads (as defined in the previous section), since all candidates in a subset code in the same phase, and therefore only very few equivalence groups need to be considered. The deconvolution is applied without regularization and for each sample separately. Further details on the deconvolution process are described in the next section. ORF candidates with an estimated activity above a threshold in at least one sample are retained.

#### ORF deconvolution

PRICE 2 estimates ORF activities by minimizing the objective function (Eq. 9), using a multiplicative update algorithm. This algorithm cannot assign negative activity values which would not be biologically plausible. Ribo-seq samples can also contain reads not generated by translating ribosomes. To account for this, PRICE 2 includes not only ORF candidates but also decoy regions. Decoy regions have an associated activity for each sample but do not consider the phase of the read or position of the read on the decoy. For each isoform with an annotated coding sequence, two decoy regions are defined: one spanning from the transcript start to the coding sequence start, and one spanning from the coding sequence start to the end of the isoform. For isoforms without an annotated coding sequence, a single decoy region spanning the whole isoform is used. The default value for the penalty strength *λ* is 10, optimized on simulated samples. Ribo-seq samples can contain positions with spuriously high read counts due to sequencing bias, which can lead to false-positive ORF calls. PRICE 2 mitigates this by applying a robust Huber regression approach. The optimization problem must be refit multiple times, but the runtime increase is modest because the previous solution is used as the initial guess for each subsequent round. The default value of the robustness parameter is 3, optimized on real data.

#### Likelihood ratio filtering

The deconvolution process retains many false-positive ORFs. We added a further filtering step to reduce the false-positive rate. For this, we optimize the unregularized objective function (Eq. 8) and compute the corresponding log-likelihood. We then greedily remove ORF candidates: starting with the lowest estimated average (over the samples) activity, each ORF candidate is removed, the activities are re-optimized, the likelihood of the reduced model is computed, and a likelihood-ratio test is used to determine whether the ORF can be removed. The number of samples is used as the degrees of freedom. The weights from the last iteration of the Huber regression are applied for the likelihood computations.

#### ORF types

Supplementary Fig. 2 is a visual depiction of the ORF types that can be assigned by PRICE 2. All transcript isoforms with the annotated cORFs from the reference are used for the ORF type assignment. The ORF types are assigned hierarchically e.g. an ORF that could be cORF on one isoform and N terminal truncated cORF on another isoform will be assigned cORF. The hierarchy is in descending order: (cORF), (N terminal extended cORF, N terminal truncated cORF),(+1 uoORF, +2 uoORF, +1 doORF, +2 doORF, +1 iORF, +2 iORF), (uORF, dORF), (pcRNA-ORF, lncRNA-ORF, varRNA-ORF). ORFs that do not fit any of the categories or two types of the same highest priority are assigned “other ORF”. Reference cORFs with a length that is not divisible by 3 are excluded, therefore giving rise to the category pcRNA-ORF: an ORF on a protein coding transcript that cannot be categorized in relation to the cORF.

#### Evaluation with simulated data

We simulated ribo-seq data to evaluate the performance of PRICE 2. We used gene locus architectures from real genes as in the reference annotation and genome. We also used coverage and cleavage models from real ribo-seq samples. We set up two simulations. In simulation one we simulated reads for 20 ribo-seq samples and 100 gene loci. In each locus we set 8 ORFs and one noise region active. One for each of the following types: cORF, uORF, +1 uoORF, dORF, +1 doORF, +1 iORF, +2 iORF, N terminal truncated cORF and the noise region. Each of the active ORFs was assigned an activity drawn from a normal distribution cORF N(2, 1), other ORFs N(1, 0.5), noise N(0.2, 0.1) clipped to 0.01. In simulation two we set 4 ORFs active in each locus (cORF, uORF, +1 iORF, +2 iORF) and a noise region. The activity for the cORF was fixed at 5 and for the noise at 0.2 and varied for the other ORF types (0.1, 0.5, 1, 2.5, 5). We simulated different numbers of runs (1, 2, 4, 8, 16, 32). Simulation 2 consisted of 30 experiments (5 activities times 6 number of runs). For each experiment 100 loci were simulated. To simulate the reads a read count was drawn from a Poisson distribution for each active region in each run according to the expected read count considering the length the activity and the run specific coverage model. Then read P-sites were distributed randomly (again considering coverage models) and upstream and downstream cleavage distances and untemplated addition status was generated according to the cleavage models. With theses characteristics the reads were mapped back to genomic coordinates.

### Creating the eval-ribo-orf-tool skill

We used Anthropic’s Opus 4.8 model with Claude code 2.1 as the Agent harness to run and evaluate ORF callers. We used the run-skill-generator skill to create another skill eval-ribo-orf-tool that, when invoked is supplied with a name of an ORF caller, which is then applied and evaluated by the agent. The eval-ribo-orf-tool was instructed to run several stages: do a web search, install the tool, prepare the input data, run the tool, convert the output to a common format and evaluate. The skill was instructed to use the 6 ribo-seq samples and, if the tool accepts them, also the 6 QTI-seq samples that were generated from the same B-cell lines [29]. PRICE 2 was applied to just the ribo-seq and to ribo-seq + QTI-seq (referred to as PRICE 2 and PRICE 2 hrt respectively). One possible bias that could lead to improved performance of our tools (PRICE 1 and PRICE 2) would be that information from the development environments leaks to the agents. We therefore strictly forbid to use any information from our system except for the input data and only run the tools as described in freely available resources on the web. In the second stage the agent installed the tool. It was instructed to preferably use a separate conda environment for each tool. The agents were instructed to fix possible problems when dependency versions were un-or misspecified and also to change code to fix compilation problems. The third stage is preparing input data. We supplied RNA-seq fastqs, QTI-seq fastq, ribo-seq fastqs, a human genome fasta, a reference GTF file and a STAR index for the genome. The agent chooses what data can be used by the tool. Importantly some tools handle both, ribo-seq and QTI-seq data. The reads of the ribo- and QTI-seq data were preprocessed: UMIs and adapters were removed and rRNA, tRNA, snRNA and snoRNA reads were removed. All the tools run on mapped reads (BAM or SAM files). Since the mapping is compute intense we instructed to keep a cache directory for the mapped files. If there is a mapping with fitting parameters in the cache the agent reuses it, else it runs a new mapping to the cache and documents the parameters. In the next step the tool is invoked. The agent was instructed to parallelize to 40 cores wherever possible or over the data samples if a single threaded task needs to apply to each data sample. It was also instructed to only use 200 GB of RAM to avoid out-of-memory errors. We designed the skill to generate reproducible workflows i.e. the agent writes bash scripts instead of directly installing or calling the tools and it documents all steps in a markdown file. If a tool runs into an error the agent investigates it and tries fixes like installing different package versions or changing few lines of code in the tool. Such changes are also documented by the agent. The tools all produce a set of ORFs that are called active but in different format. We instructed the agent to write conversion scripts for each tool. The Unified format was a tsv file with location strings in the first column with 0-based half open stop codon included intervals (e.g. 22-:22025918-22026129|22026221-22026327|22043681-22043736) and the other columns being activity or certainty like FPKM or p-values. Most of the tools run on only one data sample. In such cases the ORF sets were merged. The ORFs were translated to their protein products and Peptide-PRISM was applied. The last steps of the skill were to delete any large intermediate files (except for the mappings) and to run an analysis script to generate plots comparing all tools.

#### ORF-stock

We sought to create a comprehensive set of ncORFs. To do this we mined high quality ribo-seq samples from the public repositories Sequence Read Archive (SRA) and European Nucleotide Archive (ENA). We started with the search query:

"Homo sapiens" AND ("ribo" OR "ribosome" OR "riboseq" OR "RPF" OR "translatome" OR "footprint" OR "harringtonine" OR "lactimidomycin" OR "monosome" OR "ARTseq" OR "RiboLace")

This resulted in 9,620 run accessions, but only a subset of them were ribo-seq. To distinguish ribo-seq accessions from other protocols (mostly RNA-seq), we characterized each sample by downloading a subsample of 200,000 reads and mapping them. Each sample was labelled as ‘RIBO-SEQ’, ‘TI-SEQ’ or ‘NOT RIBO-SEQ or LOW QUALITY’ based on the fraction of reads mapping to CDS, the read length distribution of the mapped reads and the periodicity. For an example see Supplementary Fig. 7. For the exact threshold values, see Supplementary Fig. 6. The thresholds were chosen to be specific rather than sensitive for high-quality ribo-seq datasets. In total, 3,462 samples passed as ribo-seq or TI-seq. There are two caveats to these quality checks. First, most datasets are not trimmed of technical sequences; during mapping, the technical sections are soft-clipped, but they can also match the template by chance, influencing the determined footprint length and start/end position. Second, the subsampling is not done randomly, which can introduce biases in the case of position-sorted datasets. For more accurate quality checks, we computationally determined the read architecture for each sample: the type and length of the biological and technical read sections from the same read subsample that was used for the quality check. Each mapped read contains a footprint of varying length and a number of technical regions surrounding it, e.g. [5’ UMI][RT nt][footprint][3’ UMI][barcode][adapter]. More technical regions can be present, and some samples are pre-trimmed, i.e. they contain no technical regions. The idea of the read-architecture caller is that the 5’ technical regions have a fixed length, i.e. when the reads are overlaid and anchored at the 5’ end, the length and type of the regions can be determined from the entropy and mapping probability of each position 3’ of the footprint: a UMI matches only by chance ( 25%) and has high entropy, whereas a barcode also matches only by chance but has a fixed sequence, i.e. low entropy. Determining the technical regions at the 3’ end is more challenging because there is no fixed position in the reads that can be used as an anchor. Instead, the start of the adapter sequence is determined by searching for the leftmost position such that the tail of the read is a prefix of one of six adapters (illumina truseq, illumina smallrna ra3, ingolia linker, nextera, polyA, polyG). If this approach fails, the 3’-clipped sequences are searched for the most frequent 12-mer and the process is repeated. All reads are then anchored at the start of the adapter, and the 3’ technical regions are determined in the same way as for the 5’ end. Additionally, the workflow considers two technical section types that vary in length: polyA regions, which are sometimes used as adapters, and RT nucleotides/untemplated additions, which are never longer than one nucleotide but are present with a certain probability and are always A/T. For an example see Supplementary Fig. 8. The approach is not always successful in determining the read architecture, e.g. when only a very short section of the adapter is present at the 3’ end on average. In these cases our algorithm returns ‘undetermined’ instead of guessing. The 3,275 ribo-seq samples with a determined architecture were downloaded, filtered to remove contaminating reads and mapped with STAR. During the STAR mapping, the reads for each gene were counted. We kept only samples with more than 1 million mapped reads (n=3,113). We clustered the samples with Leiden clustering on the first 50 principal components of 2,000 highly variable genes. This resulted in 45 clusters that correspond to cell/tissue types. From each cluster we selected the four best samples, using the total variation distance to the uniform distribution (as a measure of periodicity) times the number of mapped reads as the quality score, and applied PRICE 2 jointly to all 180 datasets. PRICE 2 called 440,896 ORFs active. We reduced the number of ORFs by discarding those shorter than 27 nucleotides, cORFs and N-terminally truncated duplicates (n=185,433 remaining). *ORF-stock-45* was built in the same way from the best sample of each cluster (363,848 ORFs called, 160,743 remaining). For *ORF-stock-45 no RuizCuevas*, the best sample of the cluster containing the B-cell lines used for immunopeptidomics (SRR12285192) [29] was replaced by the best sample of that cluster from a different study (SRR11041307; 357,432 ORFs called, 157,770 remaining).

## Data availability

We generated no new sequencing or mass spectrometry data for this study. The ribo-seq, QTI-seq and RNA-seq data of the B-cell lines [29] are available from the Sequence Read Archive under BioProject PRJNA647736, and the matching immunopeptidomics data from PRIDE under accession PXD020620. The *ORF-stock* database and the intermediate results behind every figure and table are deposited at Zenodo (https://doi.org/10.5281/zenodo.24530344) under a CC BY 4.0 licence. *ORF-stock* can also be downloaded from https://orf-stock.erhard-lab.de. The same deposit lists the accessions of the 180 SRA/ENA runs used to build *ORF-stock* and of all 9,620 screened runs. We used the Ensembl GRCh38 annotation, release 90 for the tool benchmark and the simulations and release 116 for *ORF-stock*. The nuORFdb [40] and TransCODE [41] ORF sets are available from their original publications.

## Code availability

PRICE 2 is open source under the GNU General Public License v3.0 or later at https://github.com/kilian-m/PRICE2. RiboMine, the pipeline we used to mine, quality-check and map the public ribo-seq samples for *ORF-stock*, is at https://github.com/kilian-m/RiboMine. The Zenodo deposit (https://doi.org/10.5281/zenodo.24530344) contains the analysis and figure scripts, the ribo-seq simulator and the eval-ribo-orf-tool skill. For each benchmarked tool it also holds the install, run and conversion scripts written by the agent, together with the agent’s report. The deposit’s README gives the PRICE 2 and RiboMine commits used for each analysis.

## Acknowledgements

This research was funded by by the Deutsche Forschungsgemeinschaft (DFG, German Research Foundation) – 495215642/398367752 to FE.

