## Supplemental notes, figures and tables for "Robust ORF Deconvolution from Multiple Ribo-seq Datasets with Group LASSO Penalty"

### Supplementary Material

#### Supplementary Note 1: Evaluation on simulated ribo-seq samples

To evaluate the performance of the filtering and deconvolution steps of PRICE 2, we generated simulated ribo-seq samples. We simulated read positions and characteristics for real gene architectures using cleavage and coverage models estimated from real samples. We assigned activity to eight ORFs for each gene locus, one for each of the following types: canonical ORF (cORF), uORF, +1 uoORF, dORF, +1 doORF, +1 internal ORF (iORF), +2 iORF, and N-terminal truncated cORF (for ORF type definitions see Supplementary Fig. 2). Activity values were drawn from normal distributions with a higher expected value for cORFs than the other ORF types. We simulated data for 100 gene loci and 20 ribo-seq samples. The 100 loci contained 57,245 ORF candidates. 800 of those had non-zero activity The inactive ORF candidates were effectively filtered in multiple steps while retaining most of the active ORFs (Supplementary Fig. 3A). The first filter PRICE 2 applies on ORF candidates is based on the average in-phase read coverage with reads that can be assigned to one phase with high confidence. ORF candidates with an average (over the length) coverage below a threshold in all samples are removed. This filter removed 31,363 (54.8%) inactive ORF candidates and 7 (0.9%) active ORFs. The second filter step is a deconvolution that only considers the well fitting reads we just described and only considers groups of ORFs ending in one stop codon. This deconvolution step runs for each sample separately. This approach permits a computationally very efficient deconvolution because the equivalence groups do not distinguish by read type. This deconvolution step is not LASSO regularized. ORFs with an activity below a threshold in each sample are removed. This filter removed 14,546 (56.2%) inactive ORF candidates and 34 (4.3%) active ORFs. The next step is the main deconvolution step. Details regarding this step are described in the previous sections and in the online methods. This step was the most effective filter, removing 11,201 (98.8%) of the remaining inactive ORFs while removing 32 (4.2%) of the active ORFs. To further reduce the number of false positives PRICE 2 applies a likelihood ratio filtering step after the main deconvolution. For that, a likelihood ratio test is employed for each ORF to check whether data can also be explained without this ORF. This step removed 54 (40.0%) inactive ORF candidates and 3 (0.4%) of the remaining active ORFs. 724 of 800 active ORFs were detected and the estimated activities correlated strongly with the true activities (Supplementary Fig. 3B). The concordance correlation coefficient (CCC) of true and estimated activity values was above 0.9 for all ORF types except for N-terminally truncated ORFs (CCC=0.65) because most false negatives were N-terminally truncated cORFs. Most false negative N-terminally truncated ORFs were filtered at the deconvolution per stop codon step. These results confirm that our approach can effectively detect active ORFs. However, the accuracy depends on the ORF type, the activity, and the number of samples. To better understand what the most challenging cases are, we set up another simulation: we generated data for 100 loci with 4 active ORFs per locus, one of each: cORF, uORF, +1 iORF, and +2 iORF. We varied the activity of the non-canonical ORFs (ncORFs) and the number of samples. The cORF activity was fixed at 5. Except for some

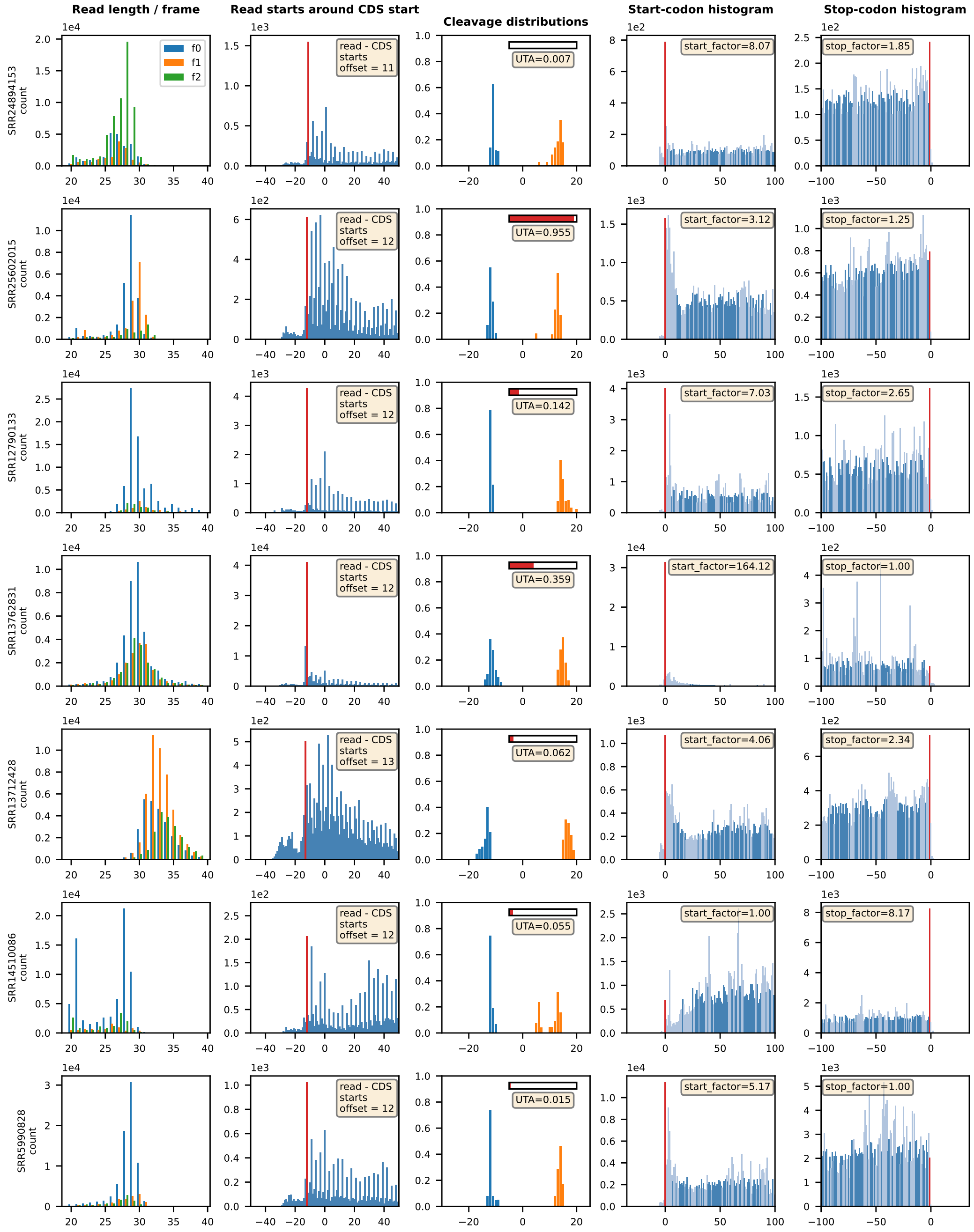


Supplementary Figure 1: Characteristics for 7 ribo-seq samples. This set was selected to show diversity in the samples. **Column 1:** Distribution of reads over different read lengths and start positions relative to the underlying reading frame. **Column 2:** Read starts around annotated coding sequence starts. Red: most frequent start to P-site shift. **Column 3:** Length distributions of cleavage probabilities upstream and downstream of ribosome P-sites. **Column 4/5:** Distribution of reads shifted to the P-site around annotated coding sequence start and stop sites. Red: start codon and one upstream of stop codon.

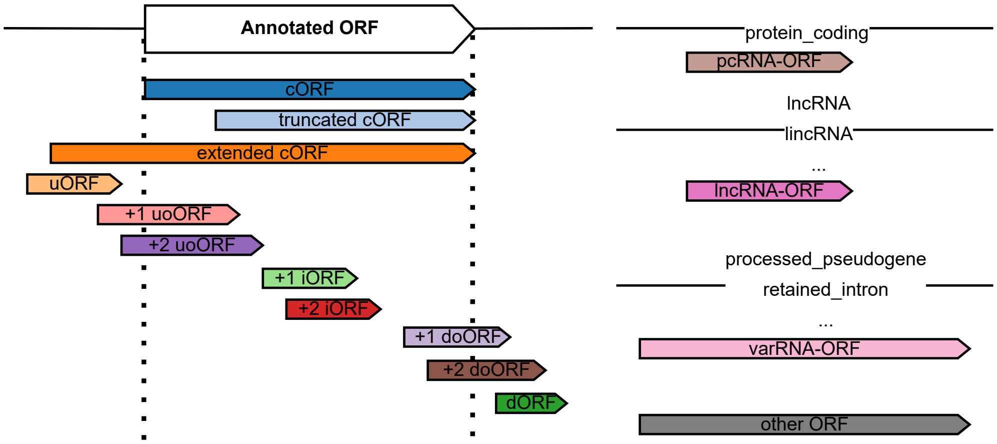


Supplementary Figure 2: ORF type definitions as used in this paper.

cases with very large ncORF activity with only a single replicate, the true positive rate (TPR) of the cORFs was above 0.9 in all simulations (Supplementary Fig. 3C). The TPR of the ncORFs increased with the number of runs and the activity (Supplementary Fig. 3D–F) and was slightly higher for uORFs compared to iORFs. Of note, even if the iORF activity was as low as 10% of the enclosing cORF activity, it could be recognized in the majority of the cases (71% or 70% for +1 and +2 iORFs, respectively), when the number of samples was large enough. In the same simulation, the number of false-positive ORFs was lowest for the simulations with only one and two sequencing runs (below 0.2 false positive ORFs per gene locus). The number of false positives was higher for more ribo-seq runs but did not show a clear correlation. The highest number of false positive ORFs was 0.34 per gene locus in 8 simulated samples at an sORF activity of 1 (Supplementary Fig. 3G). Of note, for each locus there are on average 583 ORF candidates, and in most cases zero or one were false positive, on average at estimated activity levels of 1.40.

#### Supplementary Note 2: Manual tool invocations compared to AI agent runs

We sought to compare manual tool invocations to the AI agent runs. For this we manually implemented bash scripts to apply RiboTIE [1], ORFquant [2], PRICE 1 [3] and PRICE 2 on the same data. This was done blinded to the scripts that were implemented by the AI. Post-hoc comparison of the tool invocations showed that PRICE 2 was the only tool called identically. All other tools had slightly differing routes leading to ORF sets that differed in size by up to 30%. Generally the routes taken by the AI agent seemed more reasonable but sometimes more involved e.g. the manual ORFquant call used a manually created `instructions.txt` that describes the read-length P-site offsets, the AI used RiboseQC to generate these offsets. Despite these differences between the manual and AI invocations the conclusions were identical. The consensus on the codon level was high with the largest active codon set being shared between all tools (n=14,871,609) (Supplementary Fig. 4A). PRICE 2 hrt predicted the largest set of active codons (n=28,855,855) and ORFs (n=135,438) (Supplementary Fig. 4A/B). Peptide-PRISM cryptic peptide detection found about twice as many peptides in PRICE 2 (n=91) and PRICE 2 hrt (n=101) predicted

ORFs as in the next best tool PRICE 1 (n=53) (Supplementary Fig. 4C). Only 7 peptides that were detected by any tool were not detected by PRICE 2 (Supplementary Fig. 4D). RiboTIE ORFs found more cryptic peptides in the manual evaluation because a less filtered ORF set was considered in the downstream analysis.

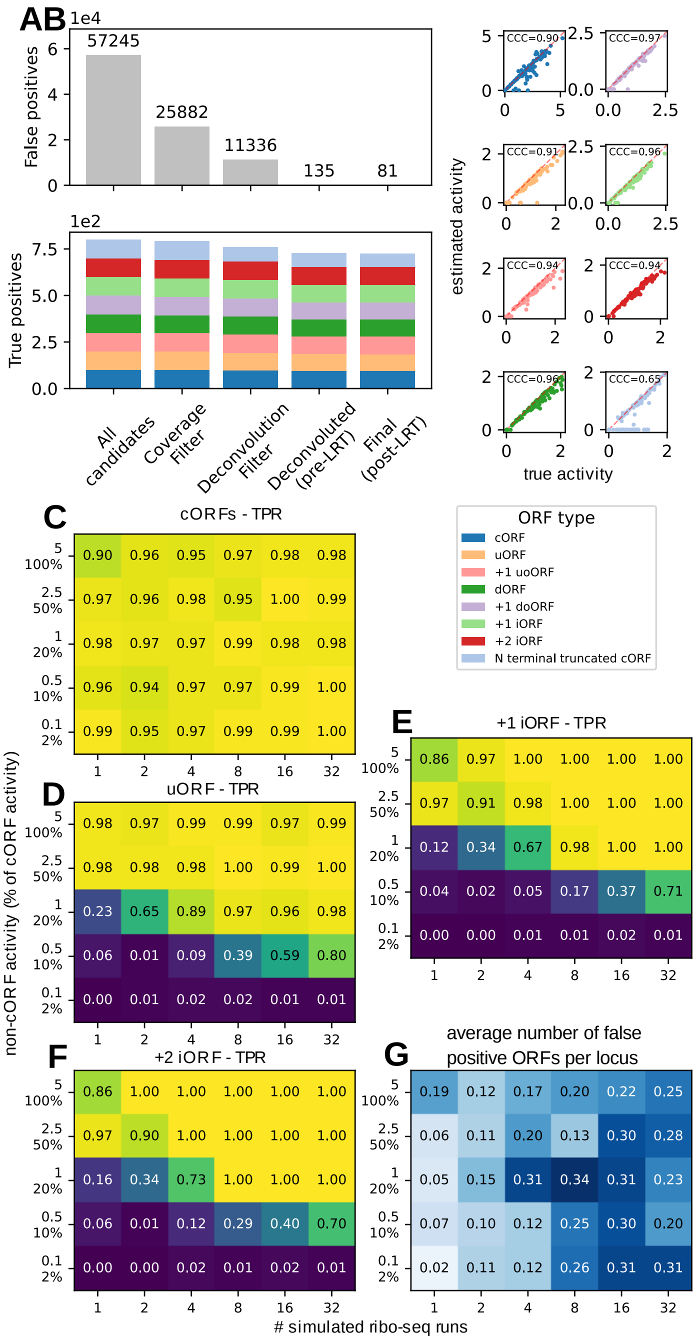


Supplementary Figure 3: Evaluation on simulated ribo-seq samples. **A**: retained false positive and true positive ORFs after each filtering step (100 loci, 20 samples, 8 active ORFs per locus (one of each type)). **B**: True and estimated ORF activities for different ORF types with concordance correlation coefficient. **E-F**: true positive rate dependent on the activity and the number of samples (tested on 100 gene loci). **G**: false positive ORFs per gene locus dependent on the activity of a +1 iORF and the number of samples. Of note, numbers in the heatmap are mean absolute frequencies and not false positive rates.

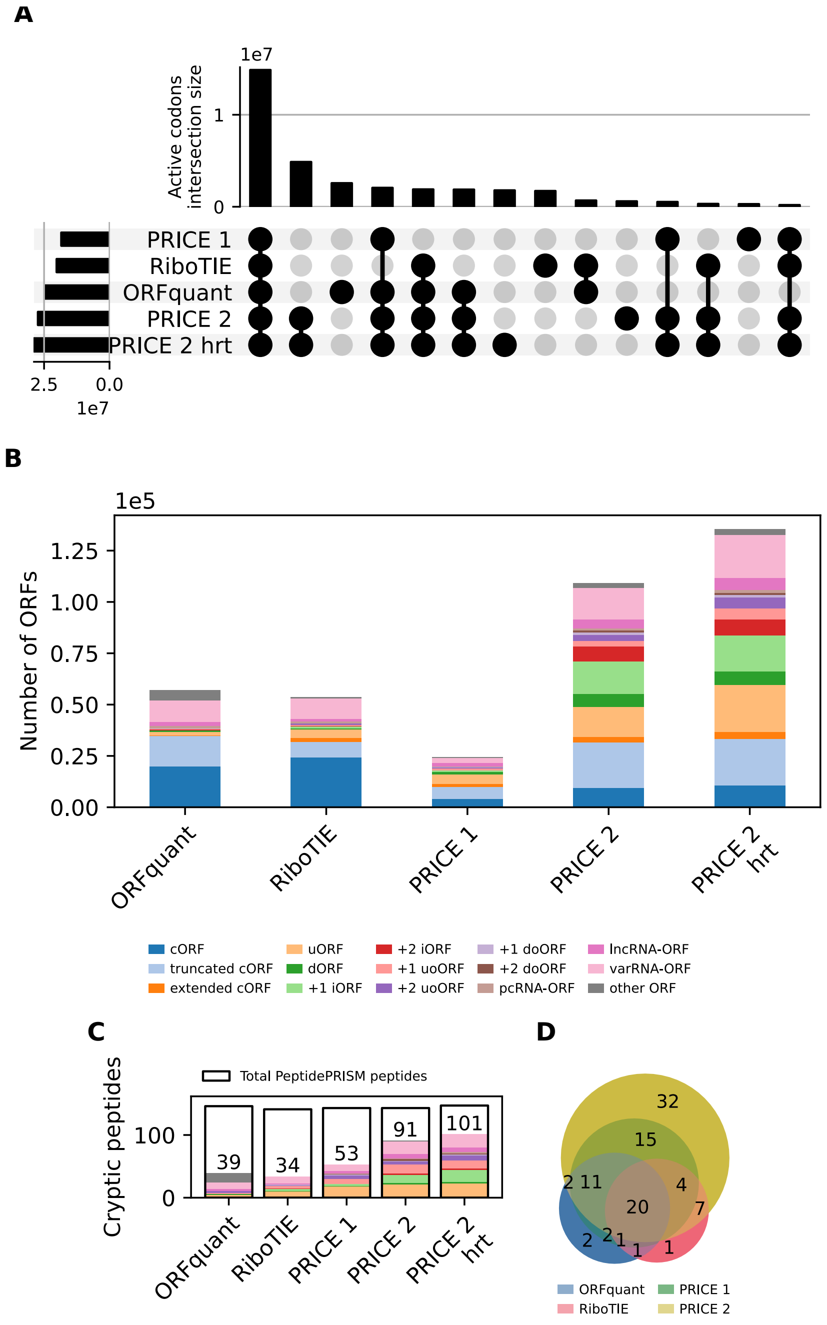


Supplementary Figure 4: Manual evaluation on real data compared to other tools. **A**: Set comparison of active codons. **B**: Number of detected ORFs by type for tools applied on the same samples. **C**: number and type of cryptic peptides that could be found in the proteins from each tool. Frames indicate totals of cryptic peptides that Peptide-PRISM detected. **D**: Euler plot of cryptic peptide sets.

Supplementary Table 1: Resource usage and ORF/cryptic-peptide detection results per tool, with implementation notes.

| Tool | Datatypes used | RAM | Runtime | ORFs | Cryptic codons | Cryptic peptides | Cryptic pep / ORFs | Cryptic pep / cryptic codons | Notes |
|---|---|---|---|---|---|---|---|---|---|
| ORF-RATER [4] | Ribo-seq, TIS | ∼20 GB | 3h 46m | 14,088 | 359,693 | 31 | 0.0022 | 8.62e-05 | py2.7; patched `get_length()` → `.length`, pinned numpy 1.16 / pandas <0.23; excluded 91 giant tfams to avoid NNLS OOM |
| ORFquant [2] | Ribo-seq | ∼73 GB group (8–23 GB/sample) | 10–28h /sample | 54,477 | 1,969,572 | 39 | 0.0007 | 1.98e-05 | `run_ORFquant` crashes at export (Bioc 3.14 seqlevels) → extracted ORFs from `_tmp_ORFquant_results`; patched BSgenome forge template |
| PRICE1 [3] | Ribo-seq | ∼168 GB | 2h 57m | 31,724 | 2,031,594 | 57 | 0.0018 | 2.81e-05 | built gedi from Maven source (no release assets); BH $q < 0.1$ == MTC count |
| PRICE2 | Ribo-seq | ∼40 GB | 37m 56s | 109,247 | 7,186,688 | 91 | 0.0008 | 1.27e-05 | built from `kilian-m/PRICE2` @ `66dfd7d` (v1.2.0), 40 workers (joint-run policy); TI-seq left unused by instruction although the tool accepts QTI-seq; RAM is 6.6 GB parent + 40 workers ≤866 MB — the 171.8 GB pipeline peak is the excluded STAR stage; no locus abandoned; v1.2.0 dropped `GenomicRegion(gi_string=...)`/`ORF_TYPES_LEVELS` → compat shims added to the shared analysis harness |
| PRICE2_hrt | Ribo-seq, TIS | ∼97 GB | 51m 30s | 135,440 | 8,575,250 | 101 | 0.0007 | 1.18e-05 | same commit, env and parameters as PRICE2 — the only difference is the 6 TI-seq (harringtonine) libraries added to the 6 Ribo-seq ones (12 datasets); +36% runtime and ×2.6 worker RSS (866 MB → 2,226 MB) vs. the Ribo-only run; the learned coverage models separate TIS (start factor 71–95) from Ribo (1.3–4.8) with no TI-seq flag; 103 Extra peptides in the run report, 101 survive the keep-mask |
| RibORF2 [5] | Ribo-seq | ∼14.1 GB group (2.5–3.2 GB/sample) | 3h 23m | 124,600 | 2,868,618 | 48 | 0.0004 | 1.67e-05 | `parameterOffset.pl` broken → reimplemented in awk; relaxed offset gate 0.6→0.5; SRR12285170 excluded (5/6 samples) |
| RiboBA [6] | Ribo-seq | ∼138 GB group (44.1–59.0 GB/sample at 3-conc.) | 1h 17m (3-conc.; 3h 23m summed) | 63,312 | 4,071,964 | 53 | 0.0008 | 1.30e-05 | R package, bias-aware EM + random forest/XGBoost; patched bioconda BSgenomeForge templates (missing `inst/extdata/` → forge fails); the tool's NCBI rRNA download returns HTTP 414 → POST-batched fallback; 6-concurrent OOM'd the 200 G cap → ran 3; output already class-thresholded, no cutoff added |
| RiboCode [7] | Ribo-seq | ≤16.3 GB/sample (∼48 GB at 4-conc.) | 3h 18m | 57,933 | 4,180,122 | 54 | 0.0009 | 1.29e-05 | metaplot `-f0_percent` 0.65 failed 4/6 → lowered to 0.5; needs STAR transcriptome BAM |
| RiboNT [8] | Ribo-seq | ∼95 GB group (17.8 GB max/sample) | 6h 49m | 82,292 | 4,908,412 | 20 | 0.0002 | 4.07e-06 | pinned scipy <1.8; stripped Ensembl `#!` header for OrfFinder parser; $O(L^2)$ Z-score slow on giant transcripts |
| RiboTIE [1] | Ribo-seq | ∼11.3 GB host (+∼12 GB GPU VRAM) | 16h 15m | 41,834 | 1,221,529 | 26 | 0.0006 | 2.13e-05 | patched upstream bitwise-`&` crash in `data.py`; cu121 torch for A10; `--max_memory` 16000 for 23 GB GPU |
| RiboTISH [9] | Ribo-seq, TIS | ∼5.3 GB group (∼0.9 GB/sample) | 1h 14m | 107,518 | 1,776,324 | 46 | 0.0004 | 2.59e-05 | needs python=3.7 (`predict` crashes ≥3.8); `--blocks` for 0-based ORF coords |
| RiboTaper [10] | Ribo-seq, RNA-seq | ∼13 GB/sample | 12h /sample (∼24.5h total) | 108,912 | 2,384,620 | 46 | 0.0004 | 1.93e-05 | reinstalled r-multitaper / r-xnomial (vendored pre-R4); offsets from metaplot peak in [-18,-6]; own BED unreliable → host-transcript mapping |

*(continued on next page)*

*(continued from previous page)*

| Tool | Datatypes used | RAM | Runtime | ORFs | Cryptic codons | Cryptic peptides | Cryptic pep / ORFs | Cryptic pep / cryptic codons | Notes |
|---|---|---|---|---|---|---|---|---|---|
| Rp-Bp [11] | Ribo-seq | ∼90 GB/sample (∼210 GB at 3-conc.) | 4–9h /sample (∼13.4h total) | 49,872 | 4,009,169 | 35 | 0.0007 | 8.73e-06 | 6-concurrent OOM'd the box → ran ≤3; systemd `MemoryMax` not enforced here; aligns internally (mapping not separable) |
| iRibo [12] | Ribo-seq | 83.4 GB | 1h 02m | 23,640 | 2,589,048 | 13 | 0.0005 | 5.02e-06 | added `#include <unordered_map>` to compile; per-sample periodicity QC would reject 2 deepest samples, but joint pooling clears it |
| ribotricer [13] | Ribo-seq | 26.1 GB group (∼4.1–4.6 GB/sample) | 3h 45m | 236,831 | 10,481,566 | 71 | 0.0003 | 6.77e-06 | single-threaded & slow (∼3.5h/sample, 6 concurrent); ∼3–5% non-mod3/no-stop ORFs are the tool's own incomplete-CDS calls (not a conversion bug) |

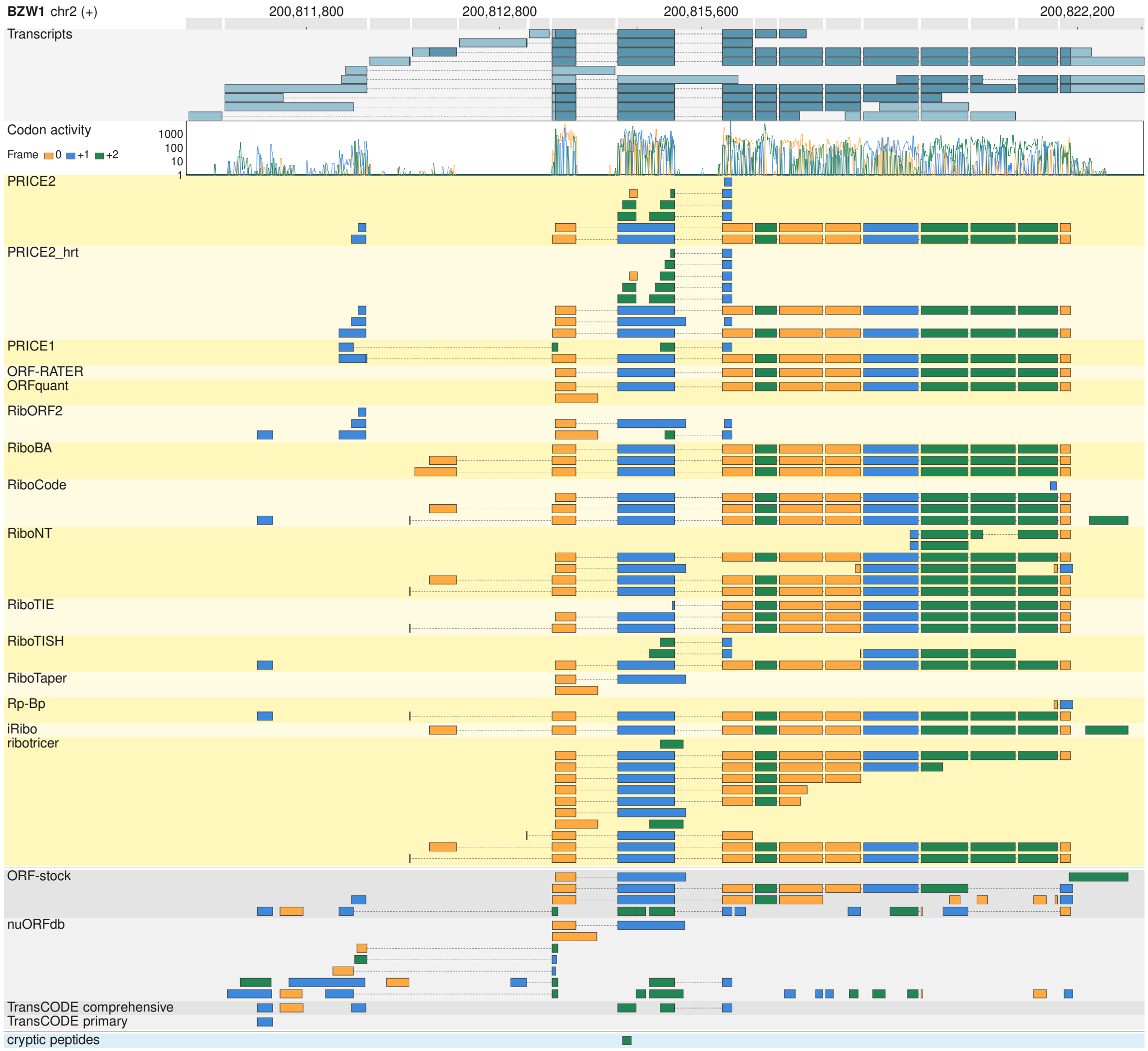


Supplementary Figure 5: ORFs detected by the tools in the BZW1 gene. Ribo-seq viewer visualization chromosome 2 positions 200,811,216 to 200,822,598. Introns in all isoforms are indicated but not shown at scale. At the top, the genomic location is shown, followed by reference transcript annotations. The codon activity track was estimated by PRICE 1. Detected ORFs for the different tools and databases are shown. Colours in the codon activity track and in the annotations from the tools signify coding phases. At the bottom the location from a peptide detected in the immunopeptidomics data is shown.

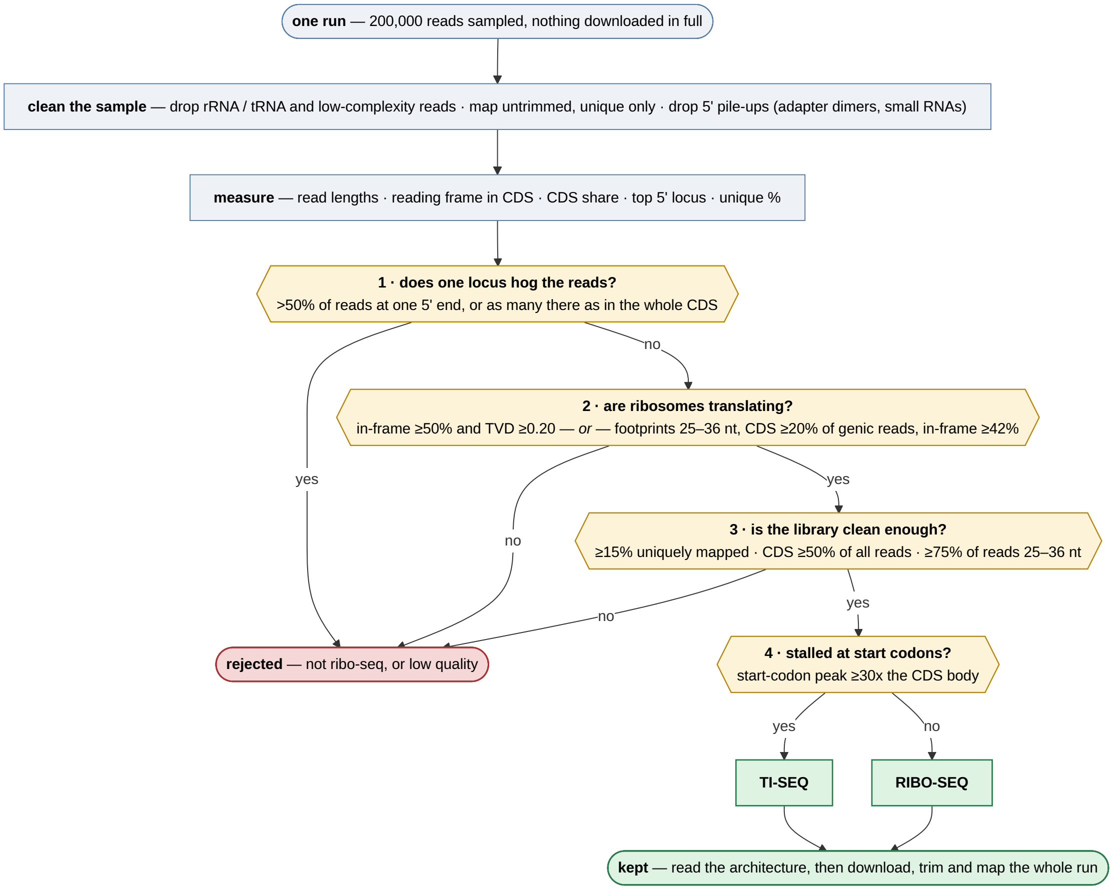


Supplementary Figure 6: Pipeline, criteria and thresholds to classify a sequencing run as ribo-seq, TI-seq or other based on a read subsample

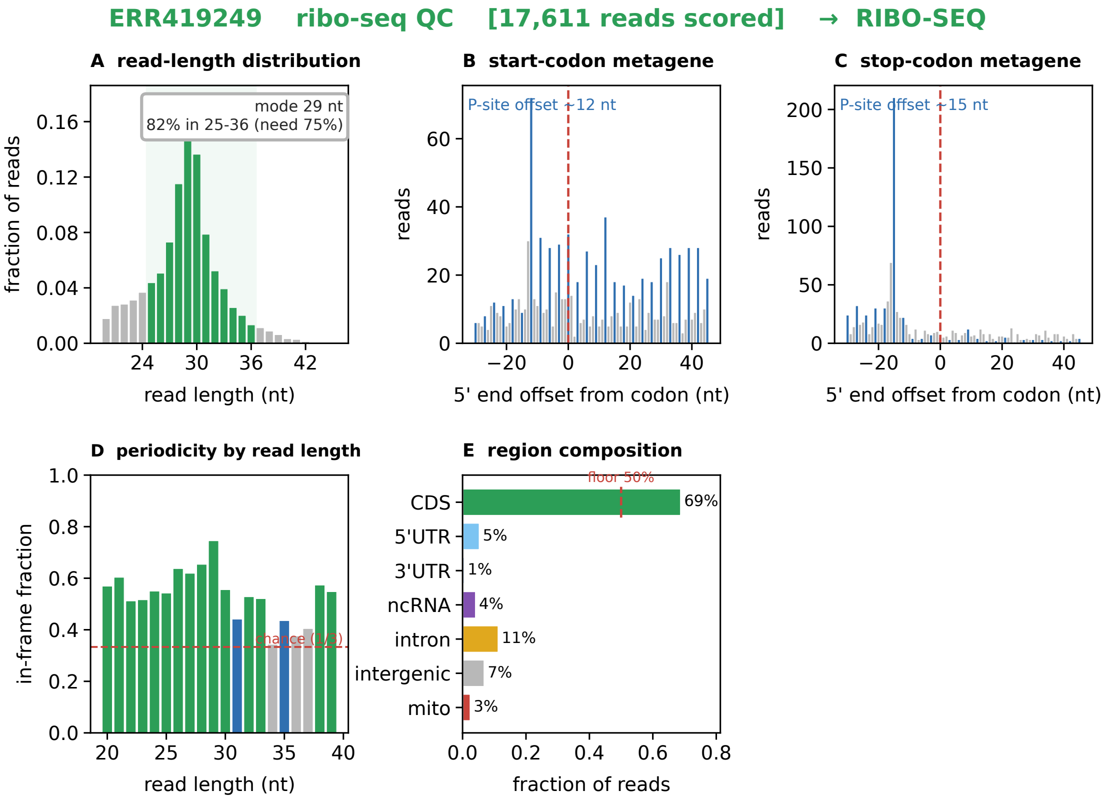


Supplementary Figure 7: Plots to check if a sequencing run is ribo-seq. **A**: read length distribution (mapping section), typical footprint lengths are highlighted. **B**: Read starts around annotated coding sequence starts. **C**: Read starts around annotated coding sequence stops. **D**: fraction of reads that start in frame for each length. **E**: fraction of reads mapping to different region types.

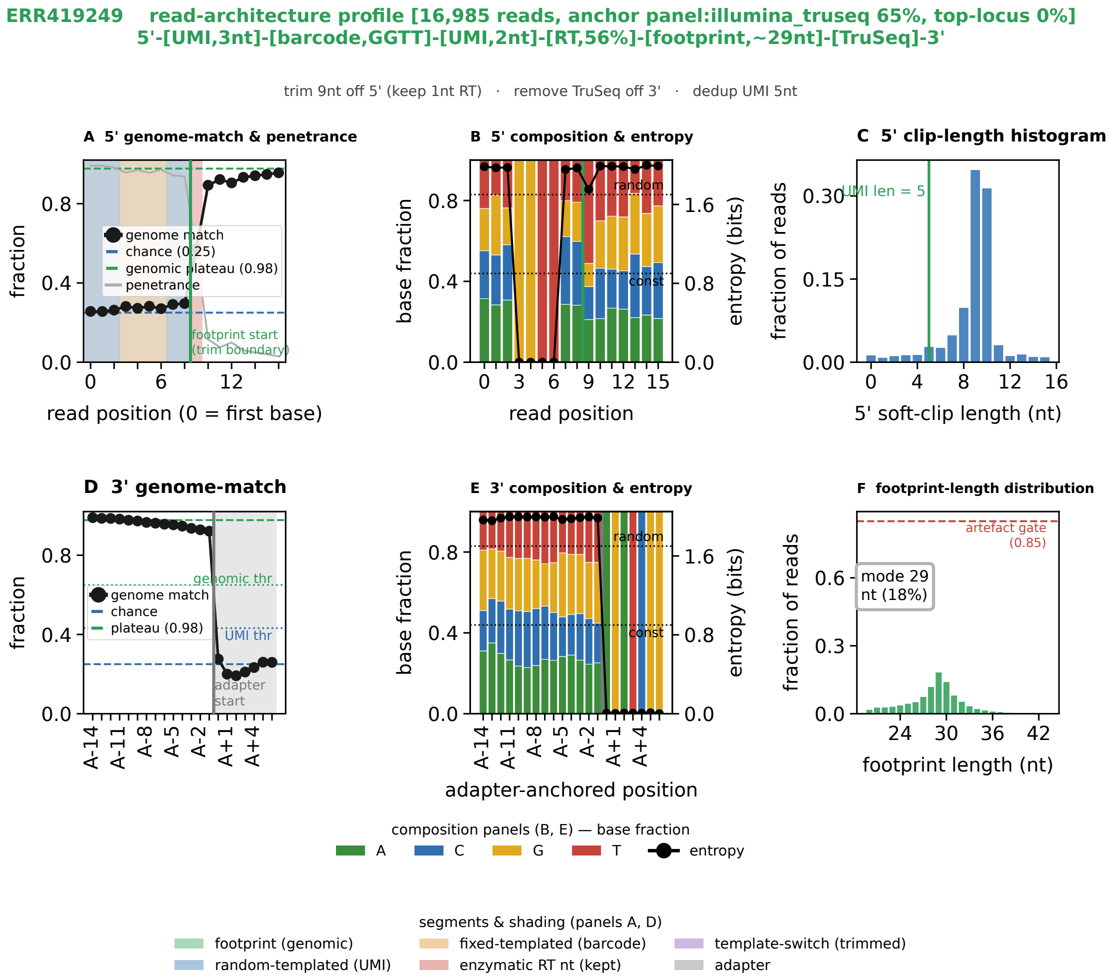


Supplementary Figure 8: Plots to infer the read architecture of a sample based on the mapping and compsition characteristics. **A**: fraction of genome matching nucleotides anchored to the 5' end of the read **B**: nucleotide composition and entropy anchored to the 5' end of the read **C**: length distribution of 5' soft-clips **D**: fraction of genome matching nucleotides anchored to the start of the adapter **E**: nucleotide composition and entropy anchored to the start of the adapter **F**: length distribution of mapping regions
